# Inhibition of mutant RAS-RAF interaction by mimicking structural and dynamic properties of phosphorylated RAS

**DOI:** 10.1101/2022.04.24.489309

**Authors:** Metehan Ilter, Ramazan Kaşmer, Farzaneh Jalalypour, Canan Atilgan, Ozan Topcu, Nihal Karakaş, Ozge Sensoy

**Affiliations:** Graduate School of Engineering and Natural Sciences, Istanbul Medipol University, Istanbul, Turkey; Cancer Research Center, Institute for Health Sciences and Technologies (SABITA), İstanbul Medipol University, İstanbul, Tukey; Medical Biology and Genetics Program, Graduate School for Health Sciences, Istanbul Medipol University, Istanbul Turkey; Faculty of Engineering and Natural Sciences, Sabanci University, Istanbul, Turkey; Department of Medical Biology, School of Medicine, Istanbul Medipol University, Istanbul, Turkey; Department of Computer Engineering, School of Engineering and Natural Sciences, Istanbul Medipol University, Istanbul, Turkey; Regenerative and Restorative Medicine Research Center (REMER), Institute for Health Sciences and Technologies (SABITA), İstanbul Medipol University, İstanbul. Turkey

## Abstract

Undruggability of RAS proteins has necessitated alternative strategies for the development of effective inhibitors. In this respect, phosphorylation has recently come into prominence as this reversible post-translational modification attenuates sensitivity of RAS towards RAF. As such, in this study, we set out to unveil the impact of phosphorylation on dynamics of HRAS^WT^ and aim to invoke similar behavior in HRAS^G12D^ mutant by means of small therapeutic molecules. To this end, we performed molecular dynamics (MD) simulations using phosphorylated HRAS and showed that phosphorylation of Y32 distorted Switch I, hence the RAS/RAF interface. Consequently, we targeted Switch I in HRAS^G12D^ by means of approved therapeutic molecules and showed that the ligands enabled detachment of Switch I from the nucleotide-binding pocket. Moreover, we demonstrated that displacement of Switch I from the nucleotide-binding pocket was energetically more favorable in the presence of the ligand. Importantly, we verified computational findings *in vitro* where HRAS^G12D^/RAF interaction was prevented by the ligand in HEK293T cells that expressed HRAS^G12D^ mutant protein. Therefore, these findings suggest that targeting Switch I, hence making Y32 accessible might open up new avenues in future drug discovery strategies that target mutant RAS proteins.

## Introduction

The RAS gene family translates into four proteins, namely HRAS, NRAS, KRAS4A, and KRAS4B, that control mitogen-activated protein kinase (MAPK), phosphatidylinositol 3-kinase (PI3K), and Ras-like (RAL) pathways ***Barbacid (1987); Malumbres and Barbacid (2003); Lu et al. (2016a); Khan et al. (2019); Duffy and Crown (2021); Simanshu et al. (2017); Ferro and Trabalzini (2010); De Luca et al. (2012); Young et al. (2013); Knight and Irving (2014***). These small G proteins act as a binary switch as the activation of the protein is modulated by two types of nucleotides, namely, guanosine-triphosphate (GTP) and guanosine-diphosphate (GDP). The exchange of GDP for GTP is maintained by guanine exchange factors (GEFs) which, in turn, activates the RAS protein ***Downward (1990); Grand and Owen (1991); Bourne et al. (1991); Wittinghofer and Pal (1991); Lowy et al. (1991); Wittinghofer and Vetter (2011); Takai et al. (2001); Lamontanara et al. (2014); Vetter and Wittinghofer (2001); Lu et al. (2016a***). Consequently, activated RAS proteins can interact with their downstream effectors, thus initiating cellular signaling pathways ***Vetter and Wittinghofer (2001); Cherfils and Zeghouf (2013); Geyer and Wittinghofer (1997); Lu et al. (2016b***). Unlike GEFs, GTPase-activating-proteins (GAPs) accelerate the intrinsic GTPase activity of RAS, which provides a control mechanism for precise termination of respective signaling pathways ***Wittinghofer et al. (1997); Lu et al. (2016a***).

RAS proteins are made up of two domains, namely, G domain (residues 1-172) and hypervariable region (173-188 or -189) ***O’Bryan (2019); Khan et al. (2020***) (Figure 1.A). The G domain consists of effector (residues 1-86) and allosteric lobes (residues 87-172). The former, which is the invariant region, harbors the P-loop (residues 10-17), Switch I (30-38), and Switch II (59-76) regions, the last two of which adopt different conformational states depending on the type of the nucleotide ***Wang et al. (2021***). In particular, Switch I/II can be found in either open or closed conformation, both of which are described depending on the position of the domain with respect to the nucleotidebinding pocket. In the open conformation, Switch I/II is far from the nucleotide-binding pocket, whereas it is closer in the closed conformation. Importantly, the former prevents effector binding while the latter favors it. Moreover, it has been also shown that Switch II becomes less stable upon effector binding, which presumably allows RAS to cycle between catalytically incompetent and competent states in a timely manner that is important for maintaining the cell homeostasis ***Johnson and Mattos (2013); Khan et al. (2020***).

**Figure 1.**
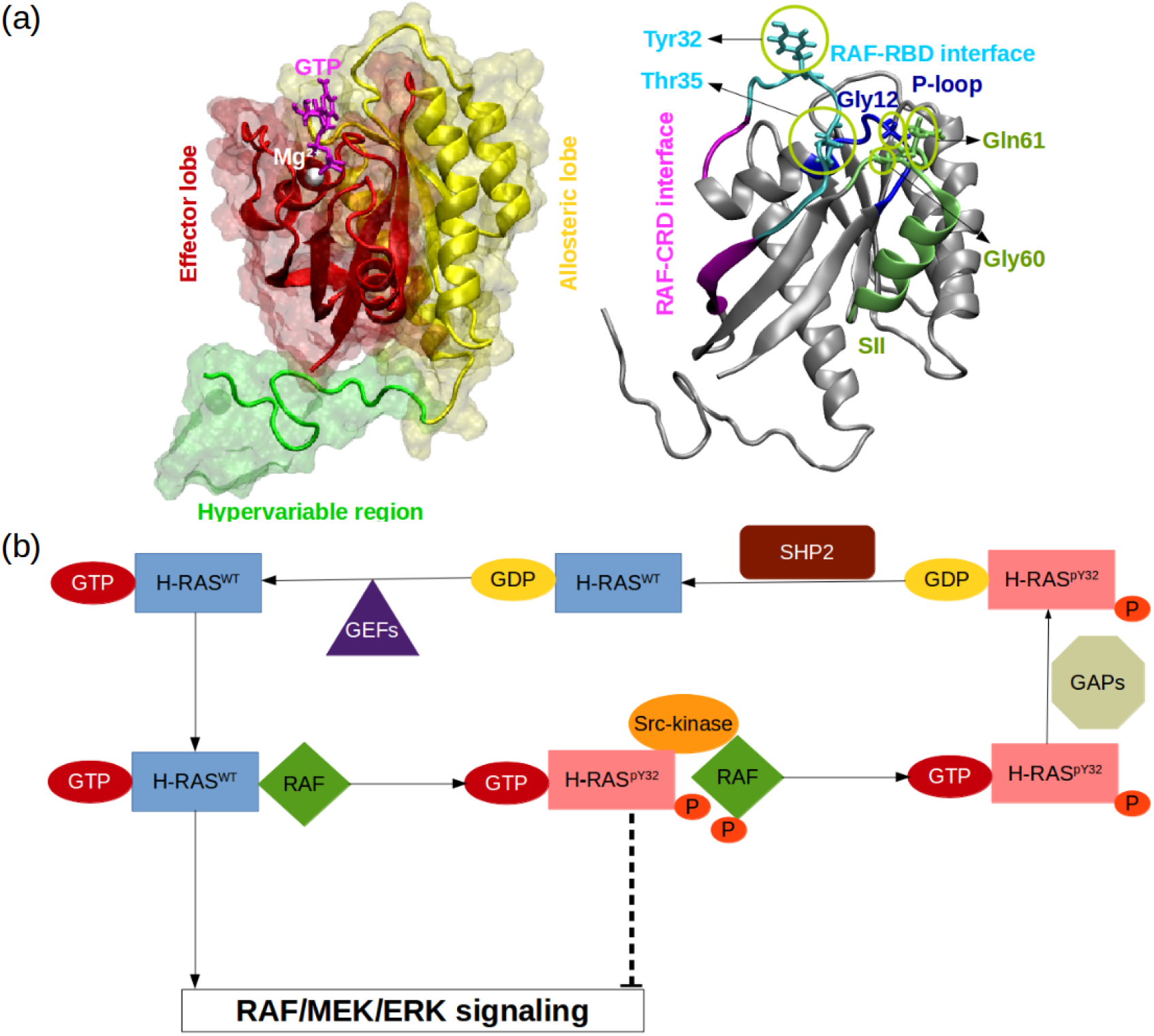
(a) Important residues/regions that play pivotal role in RAS function are shown. (b) A schematic that illustrates the impact of tyrosyl phosphorylation on the GTPase cycle of HRAS. The tyrosyl phosphorylation at the 32^nd^ position, which is mediated by Src kinase, causes impairment of RAF binding, thus terminating RAF/MEK/ERK signaling pathway as long as the phosphoryl group of Y32 is not detached by SHP2 ***Bunda et al. (2014***).

Since RAS proteins are involved in signaling pathways, which are responsible for cell growth, differentiation, and proliferation, mutations, which are frequently found at the 12^th^, 13^th^, and 61^st^ residues ***Prior et al. (2012***, 2020), cause several cancer types ***Holderfield et al. (2014); Eser et al. (2014); Prior et al. (2012); Stephen et al. (2014); McCormick (2015a***,b); ***Krens et al. (2010); Lu et al. (2016a***) as a result of attenuated GTP hydrolysis and increased nucleotide exchange rate ***Vigil et al. (2010***). For instance, HRAS^G12D^ was shown to be the dominant mutant in ductal carcinoma ***Myers et al. (2016***) caused resistance to erlotinib, which is used as an epidermal growth factor receptor tyrosine kinase inhibitor ***Hah et al. (2014***), in head and neck squamous carcinoma. As such, RAS proteins have been standing as hot targets in drug discovery studies which are focused on the development of therapeutics against cancer.

In spite of extensive efforts that have been made to develop RAS inhibitors, no molecules have yet been approved for clinical use ***Canon et al. (2019); Duffy and Crown (2021***). The undruggability of RAS proteins arises from lack of deep binding pockets on the surface of the protein and also picomolar affinity of the endogenous ligands which hinders development of competitive inhibitors ***Gysin et al. (2011); Ledford (2015); Cox et al. (2014); Milroy and Ottmann (2014***). Therefore, much attention has been focused on the discovery of allosteric sites that can regulate the function of the protein ***Buhrman et al. (2010); Ostrem et al. (2013); Fetics et al. (2015); Johnson et al. (2017); McCarthy et al. (2019); Khan et al. (2021***).

Importantly, it is well-established that the function of the protein is modulated by post-translational modifications. In particular, phosphorylation/dephosphorylation can be given as an example, which is controlled by Src-kinase and Src homology region 2 domain-containing phosphatase-2 (SHP2), respectively (Figure 1.B). It has been shown that phosphorylation of the tyrosine at the 32^nd^ position by Src-kinase attenuated RAF binding to HRAS and NRAS while elevating intrinsic GTPase activity of the proteins ***Bunda et al. (2014***) (Figure 1.B). Furthermore, recently, Kano *et al*. have implied that Src-kinase phosphorylated tyrosine residues at the 32^nd^ and 64^th^ positions of KRAS4B isoform changed conformation of Switch I and II. Consequently, this led to a decrease in intrinsic GTPase activity, thus maintaining KRAS4B in the GTP-bound state. Interestingly, phosphorylated and GTPbound KRAS4B was shown not to bind RAF, thus leaving the protein in the dark state ***Kano et al. (2019***). In the same study, it was also shown that if phosphoryl groups were removed by SHP2, then GTP-bound KRAS4B could interact with RAF and initiate signaling pathways through MAPK ***Kano et al. (2019***). Notably, it was shown that deletion or inhibition of SHP2 could slow down tumor progression, but remaining insufficient for tumor regression ***Ruess et al. (2018***). Collectively, these findings suggest that mimicking dynamics invoked by phosphorylation might provide an alternative strategy for inhibiting mutant RAS/RAF interaction.

In this study, we set out to investigate the impact of phosphorylation on the structure and dynamics of HRAS^WT^ by performing atomistic MD simulations. Comparison of the trajectory pertaining to the phosphorylated RAS with previously obtained trajectories of GTP-bound HRAS^WT^ and HRAS^G12D^ ***Ilter and Sensoy (2019***) showed that phosphorylation of Y32 increased the flexibility of both RAF-RBD and RAF-CRD (cysteine-rich domain) interfaces and pushed Switch I, in particular Y32, out of the nucleotide-binding pocket. Considering the fact that, exposed Y32 precluded RAF binding, we searched for molecules that could evoke similar rearrangements in HRAS^G12D^. To this end, we carried out virtual screening by using therapeutically-approved molecules deposited in DrugBank ***Wishart et al. (2018); Law et al. (2014); Knox et al. (2010); Wishart et al. (2008***, 2006), BindingDB ***Gilson et al. (2016); Liu et al. (2007); Chen et al. (2001b***, 2002, 2001a), DrugCentral ***Ursu et al. (2016***, 2019), and NCGC ***Huang et al. (2011***). The impact of ligands on the structure and dynamics of HRAS^G12D^ mutant was examined using molecular dynamics simulations. We showed that cerubidine, tranilast, nilotinib, and epirubicin could induce similar dynamics and structural changes which were seen in the phosphorylated RAS protein. Moreover, we also calculated the energy required for pushing Switch I out of the nucleotide-binding pocket in the absence/presence of one of the successful ligands, namely cerubidine, using perturb-scan-pull (PSP) method ***Jalalypour et al. (2020***) and showed that less energy was required for displacement of Switch I in the presence of the ligand. Importantly, we also tested the activity of cerubidine in preventing RAS/RAF interaction using immunoprecipitation assays and verified computational findings. Therefore, these results suggest that Y32 detachment from the nucleotide-binding pocket might be used as an alternative strategy for targeting mutant RAS proteins.

## Results

### Phosphorylation impacts the flexibility of RAF-RBD/RAS interface residues

The comparison of RMSF profiles showed remarkable differences in the fluctuation patterns of certain residues/domains among wild-type, phosphorylated, and mutant protein. We showed that phosphorylation increased the flexibility of Y32 as a result of repulsion between negatively charged phosphate and GTP. Interestingly, we also observed that post-translational modification increased the flexibility of the residues that are involved in the RAF-RBD/CRD interaction interface as shown in Table 1. The RAF-CRD has been shown to play an important role in anchoring RAF to the membrane and enhancing RAS-RAF interaction ***Travers et al. (2018***) by binding G60 and Q64 residues of RAS as revealed by NMR and mutagenesis studies ***Drugan et al. (1996***). Interestingly, we observed that flexibility of G60 increased upon phosphorylation (See Table 1), hence presumably interfering interaction of RAS with RAF-CRD. Moreover, phosphorylation also increased flexibility of Q61, which might impact GAP-mediated GTPase activity of the protein as GAP stabilizes the catalytically-competent conformation of catalytic residue Q61 ***Simanshu et al. (2017***). Of note, the flexibility of RAF-RDB interface residues were higher in HRAS^G12D^ than in HRAS^WT^.

**Table 1.** The backbone RMSF values of key regions/residues pertaining to HRAS^WT^, HRAS^pY32^, and HRAS^G12D^.

| Residue/Region-RMSF (Å) | HRAS<sup>WT</sup> | HRAS<sup>pY32</sup> | HRAS<sup>G12D</sup> |
| --- | --- | --- | --- |
| Y32 | 1.0 | 1.8 | 1.4 |
| RAF-RBD interface residues | 1.1 | 1.5 | 1.9 |
| RAF-CRD interface residues | 0.8 | 1.2 | 0.8 |
| G60 | 1.9 | 3.4 | 1.8 |
| Q61 | 2.2 | 3.6 | 2.4 |

### Phosphorylation pushes Switch I and Y32 out of the nucleotide-binding pocket of RAS

As shown in Table 1, the flexibility of residues, which interact with RAF-CRD domain, increased upon phosphorylation. Since these residues surround the Switch I domain, we sought to investigate whether the opening of the nucleotide-binding pocket was impacted by measuring the distance between C*α* atoms of the G/D12 and P34 residues throughout the trajectories. We showed that phosphorylation pushed Switch I out of the binding pocket as the distance between G12 and P34 residues increased compared to wild-type and mutant protein (Figure 2.A). Consequently, this makes the nucleotide-binding pocket more accessible to waters, as evident from the number of waters measured within 5 Å distance of GTP: 90.7±0.1, 103.4±0.1, and 88.9±0.1 for HRAS ^WT^, HRAS^pY32^, and HRAS^G12D^, respectively, thus, presumably, modulating intrinsic GTPase activity of the protein ***Bunda et al. (2014***). Interestingly, the nucleotide-binding pocket could adopt more open conformations in HRAS^G12D^, yet as not frequent as seen in HRAS^pY32^.

**Figure 2.**
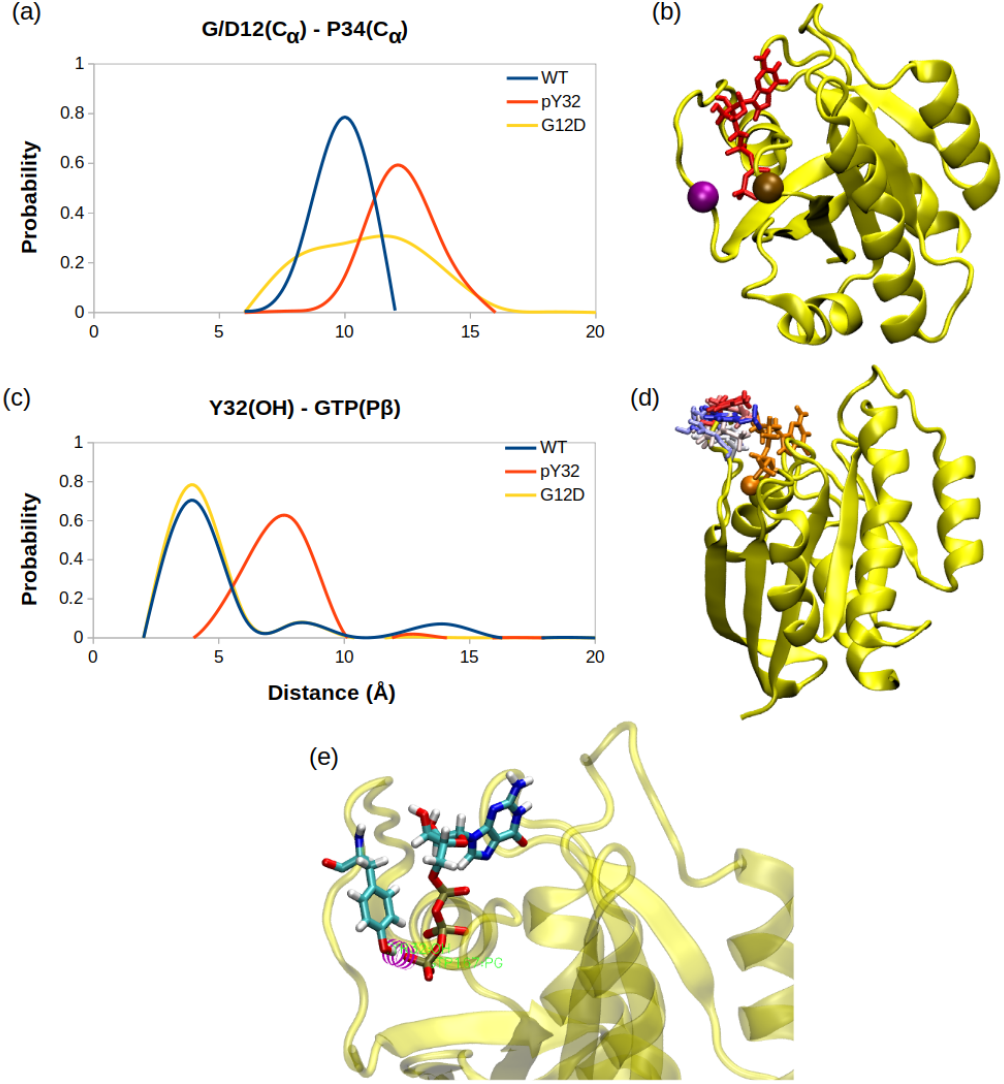
(a). The probability distribution of the distance measured between the C*α* atom of 12^th^ and 34^th^ residues. (b) C*α* atoms of 12^th^ and 34^th^ residues are shown on the crystal structure of HRAS^WT^ (PDB ID: 5P21) in vdW representation and colored with ocher and purple, respectively, whereas GTP is shown in the licorice and colored with red. (c). The probability distribution of the distance measured between the side-chain oxygen atom of Y32 and P*γ* of GTP. (d) The orientational dynamics of Y32 in the HRAS^pY32^ trajectory. (e) The H-bond formed between side-chain of Y32 and P*γ* of GTP in HRAS^G12D^ is shown in purple.

Having observed phosphorylation-induced modulation in the flexibility of Y32, we also examined the positioning of the residue by measuring the distance between the side-chain oxygen of Y32 and P*γ* atom of GTP. We showed that Y32 formed a hydrogen bond with the P*γ* atom of GTP in both HRAS^WT^ and HRAS^G12D^ which stabilized the residue in the vicinity of the nucleotide-binding pocket (Figure 2.C, and E.). However, the hydrogen bond was not formed in HRAS^pY32^, thus repositioning Y32 far from the pocket, thus making it exposed to the environment, as evidenced by relatively longer distances measured (Figure 2.C). In addition to the position, we also explored orientational preference of Y32 with respect to the nucleotide-binding pocket by measuring dihedral angles pertaining to backbone and side-chains of Y32, namely *ϕ*/*ψ* and *χ*_1_/*χ*_2_ angles. There was no remarkable difference in backbone dihedrals and *χ*_1_, whereas *χ*_2_ angle distribution was different among the systems studied. Specifically, Y32 displayed two peaks at -100 --90^°^ and 80-90^°^ in the phosphorylated RAS, whereas it adopted 60-70^°^ in the mutant and wild-type HRAS (Figure 3.A). It is important to point that Y32 adopted 80^°^ in the crystal structure of allosteric inhibitor-bound KRAS4B^G12D^ (PDB ID:6WGN) ***Zhang et al. (2020***), where the residue was exposed and far from the nucleotide-binding pocket as the distance measured between the side-chain of Y32 and P*γ* atom of GTP was 16 Å.

**Figure 3.**
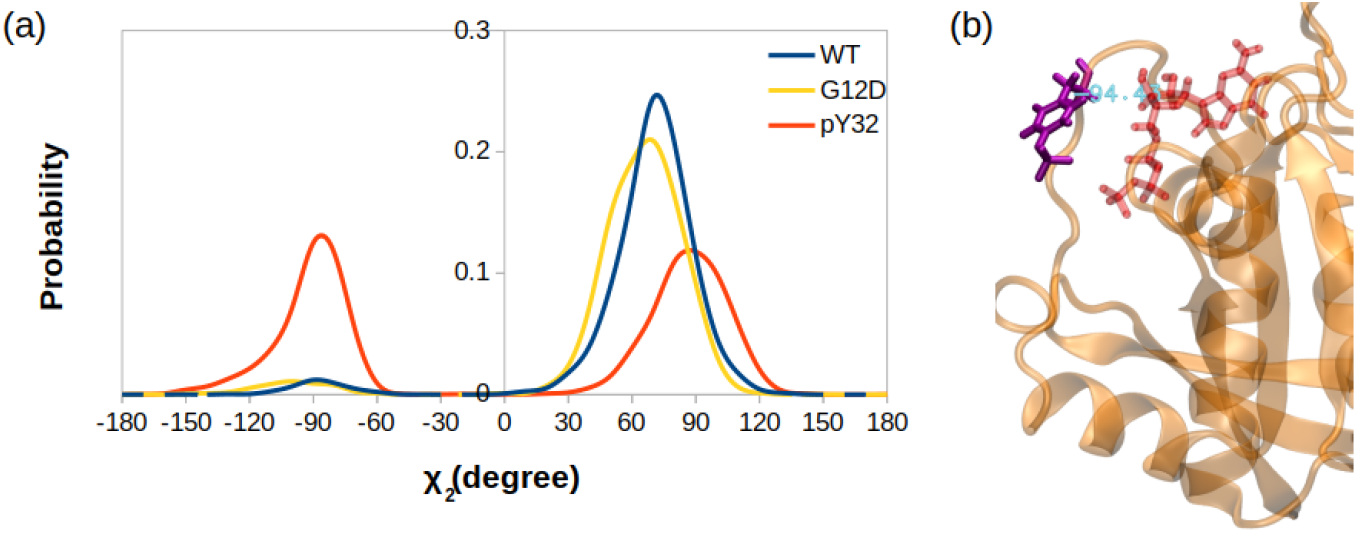
(a) The probability distribution of the measured *χ*_2_ angles of HRAS^WT^, HRAS^G12D^, and HRAS^pY32^. (b) A representative exposed state of Y32 obtained from the trajectory of the phosphorylated system.

Herein, it is important to mention that exposed conformation of Y32 was not observed in the trajectories pertaining to RAF-RBD-bound HRAS^WT^ as shown in our earlier study ***Ilter and Sensoy (2019***). Therefore, this finding suggests that exposure of Y32 might occlude the interaction interface formed between RAS and RAF-RBD and exposure of Y32 might facilitate water attacks to P*γ* of GTP, hence increasing intrinsic GTPase activity, in accordance with experimental data ***Bunda et al. (2014***).

### Global dynamics reveals a possible binding site near the nucleotide-binding pocket in HRAS^G12D^

Besides local analysis, the collective dynamic properties of the systems were also explored by calculating the principal components of their global motions. To do so, trajectories of HRAS^WT^, HRAS^G12D^, and HRAS^pY32^ were projected along their first three eigenvectors, which reflect *ca*. more than 50% of the overall dynamics, and compared to each other to investigate global conformational rearrangements induced by phosphorylation and mutation. Consequently, in line with the RMSF profiles, it was shown that G12D mutation significantly altered dynamics of Switch I domain, in particular, the RAF-RBD interaction interface. Although RAF-RBD interface dominated the collective motion in the mutant compared to phosphorylated HRAS, contribution of Y32 to the first two eigenvectors was higher in HRAS^pY32^ (1.74) than in wild-type (0.43) and mutant protein (0.95) (Figure 4.A & .B).

**Figure 4.**
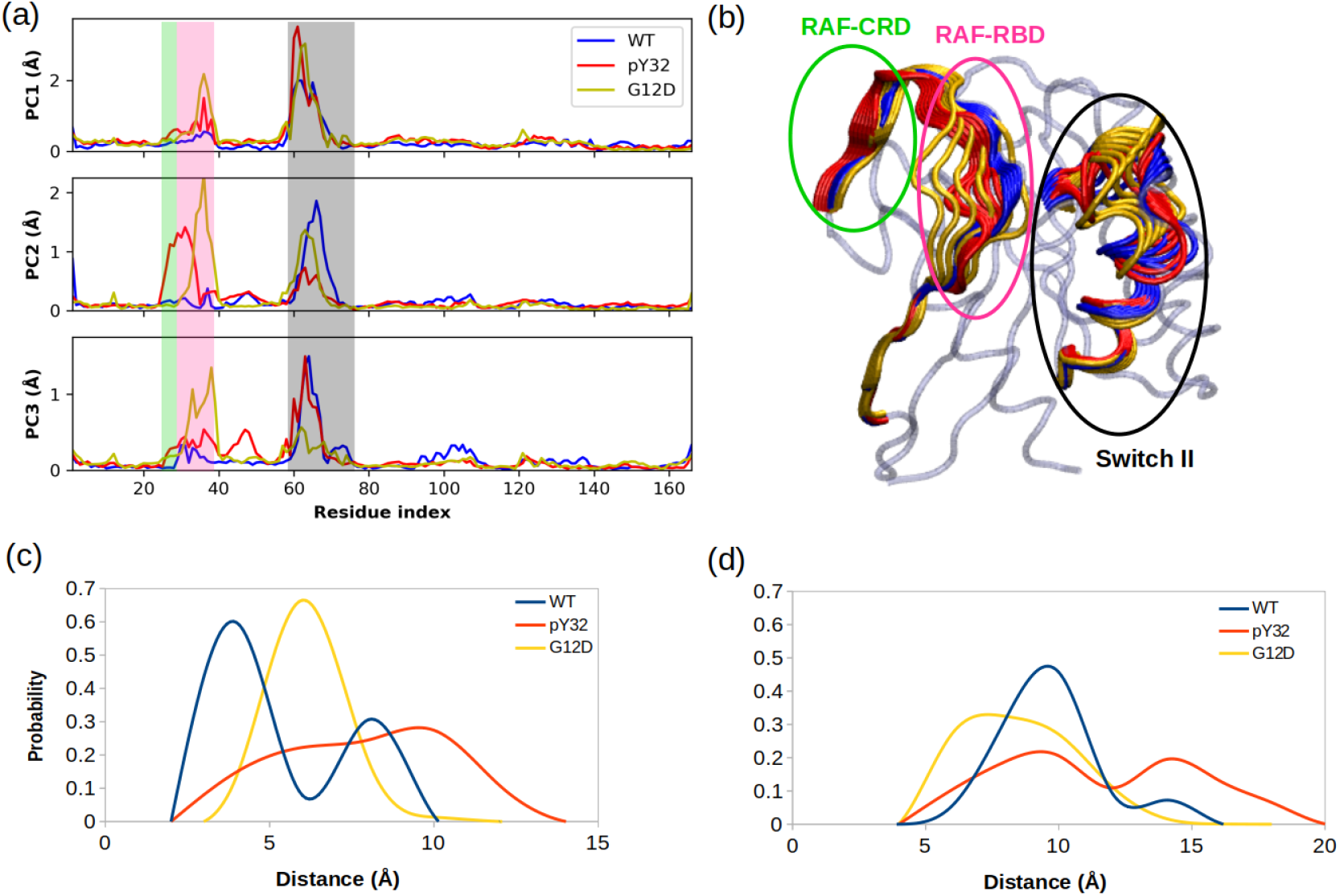
(a) Fluctuation of C_*α*_ atoms pertaining to HRAS^WT^, HRAS^G12D^, and HRAS^pY32^ along the first three eigenvectors. The RAF-CRD & -RBD interaction interfaces, as well as Switch II, are shaded in the green, pink, and black rectangles, respectively. The eigen RMSF of Y32 pertaining to the phosphorylation system is pointed out by a dark violate bead. (b) The projected trajectories of the systems studied along with the first principal component, where the thickness of the ribbons are correlated the contribution of domain to the collective dynamics. The probability distribution of the distance between (c) the backbone amide of G60 and P*γ* of GTP is shown, and (d) the side-chain oxygen of Q61 and P*γ* of GTP is shown.

Interestingly, the contribution of Switch II, which harbors both G60 and Q61, to the overall dynamics was similar in HRAS^pY32^ and mutant protein (Figure 4.A). However, G60 and Q61 were positioned closer to the nucleotide-binding pocket in the mutant HRAS than in phosphorylated HRAS (Figure 4.C and D).

Having observed higher flexibility at the RAF-RBD interface in the mutant, we set out to investigate if the site can be considered as a possible binding pocket that can accommodate small molecules to modulate the dynamics of Switch I. To do so, we clustered the trajectory of the mutant HRAS by considering probability distributions of distances between the (i) side-chain oxygen of T35 and P*γ* of GTP (ii) backbone amide of G60 and P*γ* of GTP, and (iii) side-chain oxygen atom of Q61 and P*γ* of GTP of HRAS^G12D^, which represent different conformational states of the nucleotidebinding pocket, according to the structural studies ***Vetter and Wittinghofer (2001); Shima et al. (2010); Araki et al. (2011); Pai et al. (1990); Huang et al. (1998); Buhrman et al. (2010***). There were three states described for T35, labelled as state 1, 2, and 3, each of which sampled distances in the range of 3.0-5.0 Å 6.0-9.0 Å and 12.0-16.0 Å respectively (Figure 5.A). Similarly, G60 could also adopt three states, namely state 1, 2, and 3, which corresponds to distance range between 5.0-7.0 Å 2.0-4.0 Å and 8.0-9.0 Å respectively (Figure 5.B). Moreover, Q61 could sample distances in the range of 8.0-10.0 Å 4.0-7.0 Å and 10.0-14.0 Å so adopting three states, namely state 1, 2, and 3 (Figure 5.C). In light of clustered conformations, the most probable conformation that adopts values pertaining to State 1 in each atom-pair distances was picked up from the trajectory of HRAS^G12D^. The possible binding pockets on the surface of mutant HRAS were identified and evaluated by comparing SiteMap scores. Eventually, the pocket, which had relatively higher SiteScore, enclosure and lower exposure, was selected to be used further (Table 2.). The binding pocket, which was identified on the selected conformation, was next to Switch I. Considering the fact that this domain i) includes residues that mediate RAF binding, ii) acts as a regulator for intrinsic GTPase activity, and iii) dominates the collective dynamics of the mutant protein, the region was used as the target binding pocket in the subsequent steps of the study.

**Table 2.** The SiteMap scores of possible pockets found on the surface of the most probable conformation of HRAS^G12D^.

| SiteScore | Size | DScore | Volume | Exposure | Enclosure | Contact | Phobic | Philic | Balance | Don/acc |
| --- | --- | --- | --- | --- | --- | --- | --- | --- | --- | --- |
| 1.028 | 194 | 0.919 | 466.140 | 0.478 | 0.740 | 1.010 | 0.259 | 1.425 | 0.182 | 0.932 |
| 0.701 | 25 | 0.668 | 81.290 | 0.632 | 0.691 | 1.059 | 1.474 | 0.664 | 2.219 | 0.915 |
| 0.656 | 22 | 0.629 | 101.870 | 0.776 | 0.638 | 0.842 | 0.959 | 0.596 | 1.609 | 12.216 |

**Figure 5.**
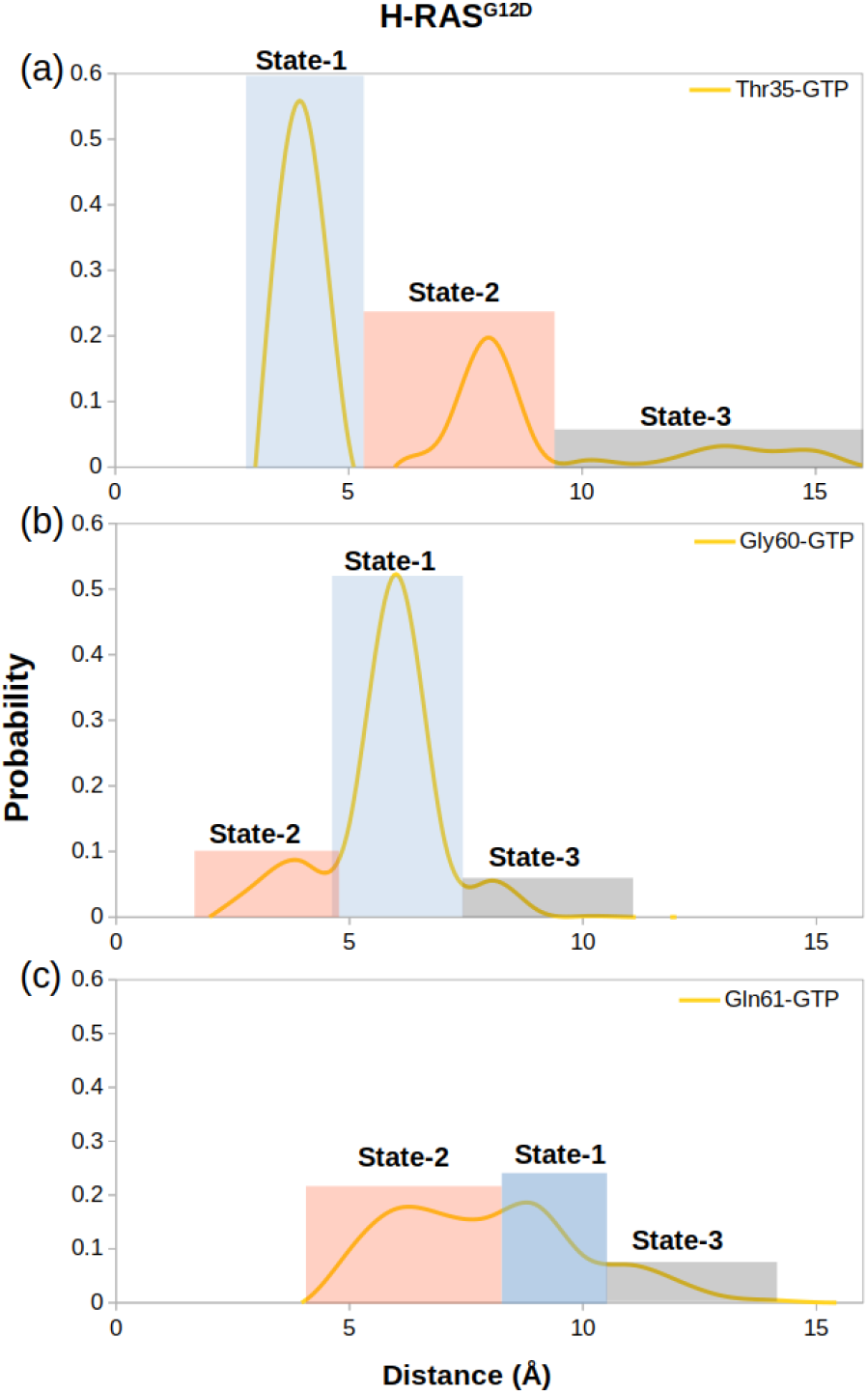
The probability distribution of the measured distance between (a) the side-chain oxygen atom of T35 and P*γ* of GTP, (b) the backbone amide of G60 and P*γ* of GTP, and (c) the side-chain oxygen atom of Q61 and P*γ*, where the sampling range for calculating the frequency of each interval was adjusted as 1 Å.

### Small therapeutic molecules distort the RAF binding interface and pushes Y32 out of the pocket

The pharmacophore groups of the binding site identified on the surface of HRAS^G12D^ were modeled with respect to both geometrical and chemical properties of residues 29-34. DrugBank ***Wishart et al. (2018); Law et al. (2014); Knox et al. (2010); Wishart et al. (2008***, 2006), DrugCentral ***Ursu et al. (2016***, 2019), BindingDB ***Gilson et al. (2016); Liu et al. (2007); Chen et al. (2001b***, 2002, 2001a), and NCGC***Huang et al. (2011***) databases were searched for molecules that could contain at least 3 features of the modeled pharmacophores and have molecular weight lower than 550 kDa. A total of 4292 molecules was retrieved from the databases (Figure 6.A). Then, these molecules were docked to the identified binding pocket on the surface of HRAS^G12D^ and ligands were evaluated with respect to their spatial organization around the nucleotide-binding pocket and GScores, which is a term that is used to score binding poses in Schrodinger. Considering the close interaction observed between Y32 and GTP in HRAS^G12D^, we prioritized the ligands, which disrupted interaction between the nucleotide and Y32. The impact of four ligands satisfying this criterion, namely cerubidine, tranilast, nilotinib, and epirubicin, (Figure 6.B, .C, .D, & .E) was further tested by performing MD simulations using the ligand-HRAS^G12D^ complex (See Table S2 for respective simulation times).

**Figure 6.**
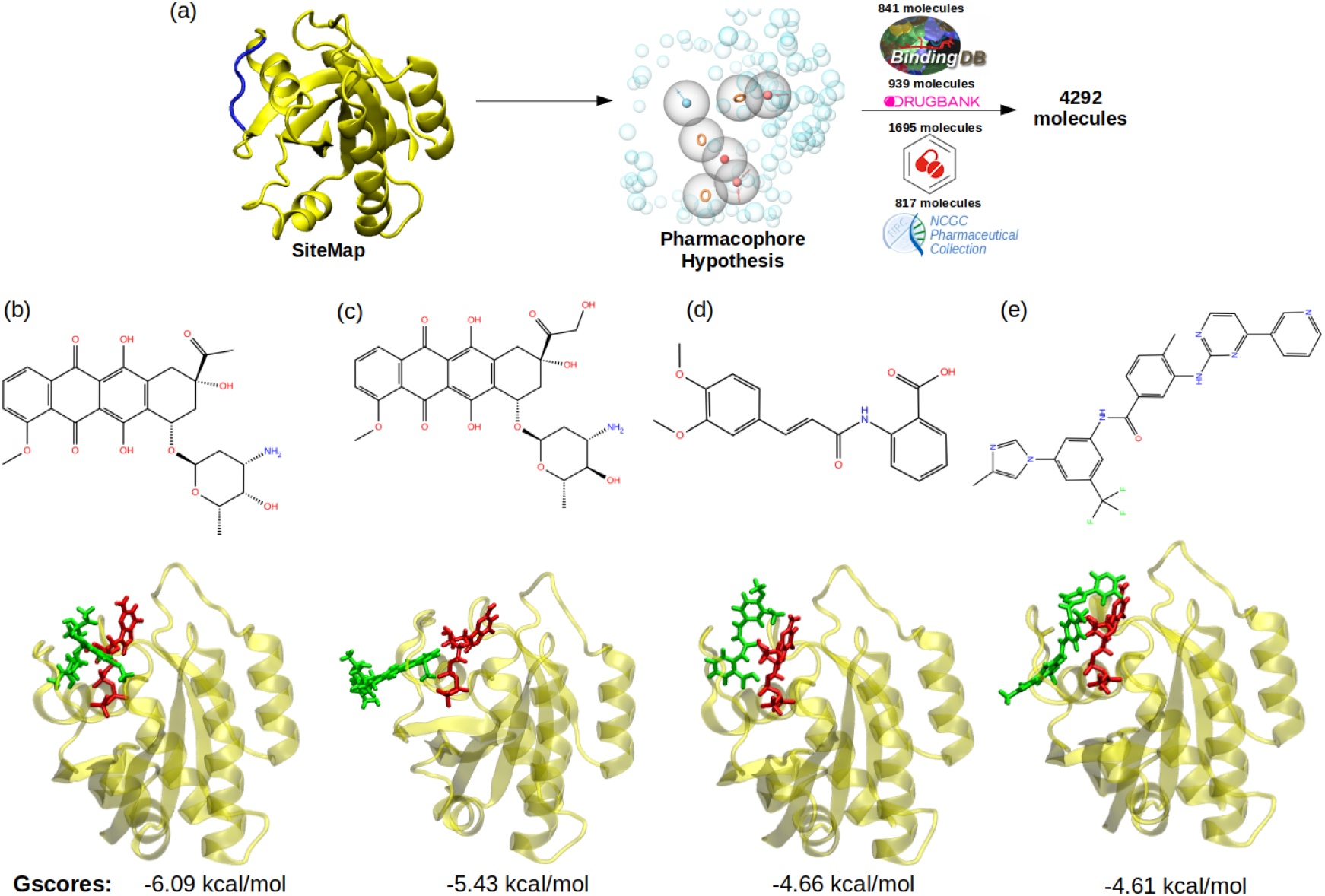
(a) A schematic that summarizes the virtual screening workflow done for the identified binding pocket on the most frequently sampled conformation of HRAS^G12D^. The 3D structures and corresponding GScores of (b) cerubidine, (c) epirubicin, (d) tranilast, and (e) nilotinib are shown. GTP is shown in licorice and red.

The ligand-HRAS^G12D^ trajectories were analyzed based on the fluctuation pattern of RAF-RBD, RAF-CRD, and Y32. Moreover, the distances measured between G/D12 and P34 as well as Y32 and GTP were also compared to those of HRAS^G12D^ and HRAS^pY32^. In that way, the capability of the ligands in distorting Switch I domain, widening the nucleotide-binding pocket, and displacing Y32 could be investigated. Accordingly, ligands, which could (i) increase the flexibility of RAF-RBD and -CRD interfaces, and (ii) displace Switch I and Y32 from the nucleotide-binding pocket were considered successful in terms of preventing HRAS^G12D^/RAF interaction. We showed that all the ligands, namely cerubidine, nilotinib, tranilast, and epirubicin, considerably increased the flexibility of the RAF-RBD interaction interface (See Table S2) than in HRAS^G12D^. Moreover, the flexibility of Y32, also significantly increased by all the ligands, except nilotinib ***Bunda et al. (2014); Kano et al. (2019***) (Table S2).

We also examined the wideness of the nucleotide-binding pocket and the positioning of Y32 by measuring the distances between G/D12, respectively. We showed that cerubudine was more likely to trigger displacement of Switch I and Y32 away from the nucleotide-binding pocket, whereas the impact of nilotinib and epirubicin was not remarkable (See Figure 7.A and .B). This, in turn, explains accommodation of relatively higher number of waters within the nucleotide-binding pocket in cerubidine-bound HRAS^G12D^ (Table S1). Therefore, it is tempting to suggest that cerubidine can help elevate the intrinsic GTPase activity of the mutant RAS by exposing GTP to water.

**Figure 7.**
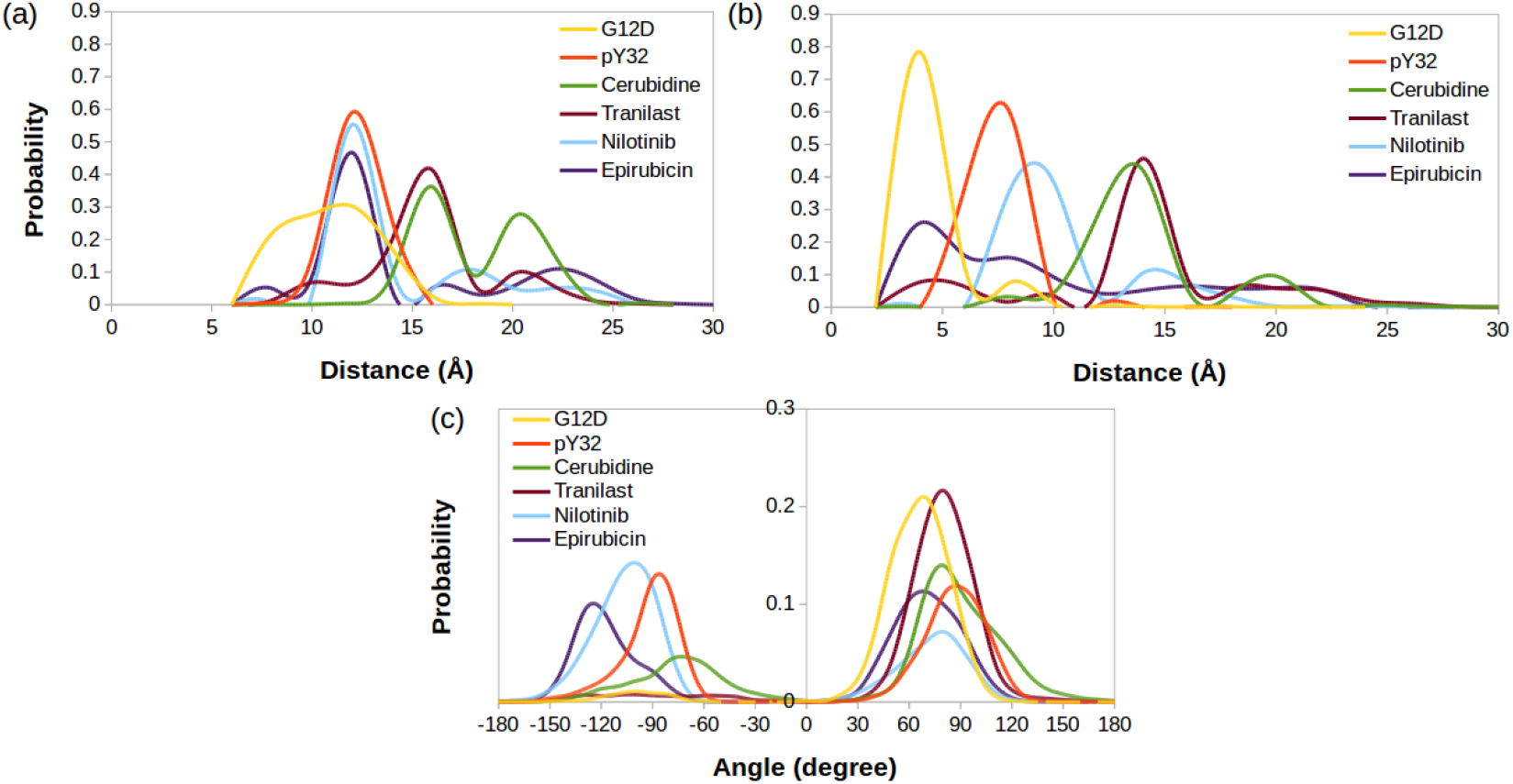
The probability distribution of (a) the distance between C_*α*_ atoms of G/D12 and P34, (b) the distance between the side-chain oxygen atom of Y32 and P*γ* of GTP, (c) *χ*_2_ in HRAS^G12D^, HRAS^pY32^, and cerubidine-, tranilast-, nilotinib-, and epirubicin-bound HRAS^G12D^

Also, Y32 adopted similar *χ*_2_ angle in cerubidine-bound HRAS^G12D^ to that in the phosphorylated RAS (Figure 7.C). Considering similarities between ligand-bound HRAS^G12D^ and HRAS^pY32^, cerubidine can be thought to have relatively more potential for preventing HRAS^G12D^/RAF interaction. Therefore, we used cerubidine in the subsequent steps of the study to test our proof-of-concept.

### Perturb-Scan-Pull method reveals that displacement of Switch I and Y32 in HRAS^G12D^ is favored in the presence of cerubidine

We further set out to investigate if displacement of Switch I and Y32 is energetically favorable in the presence of the cerubidine. To do so, we applied perturb-scan-pull (PSP) method ***Jalalypour et al. (2020***), which was developed to investigate conformational transitions in proteins, on cerubidinebound HRAS^G12D^. In this approach, initial and target states are described and the most possible path for transitioning between the initial and the target state is determined by calculating the overlap between the states. The maximum overlap is thought to give the optimum conformational transition path. To be consistent with the previous analyses, we used the same reaction coordinates, such as the distance between i) C_*α*_ atoms of D12 and P34, and ii) the backbone amide of G60 and P*γ* of GTP, as the reaction coordinates, which, reflected dynamics of Switch I and II, respectively. Accordingly, we described three and two states for Switch I and II, respectively, considering the conformations obtained by clustering of HRAS^G12D^ trajectory. Accordingly, for Switch I, if the measured distance between C*α* atoms of D12 and P34 is less than 8 Å, Switch I is grouped as in the closed state. On the other hand, when the atom-pair distance is above 16 Å, Switch I grouped as in the open state. The distance between 8 and 16 Å is grouped as the partially open state of Switch I. Likewise, we also determined the state of the Switch II by measuring the distance between the backbone amide of G60 and P*γ* of GTP. If the distance is above 11 Å, Switch II is grouped as in the open state, if not, in the closed state. In light of these atom-pair distances, the initial state was described as the closed state of both Switch I and II domains, since it was the most frequently sampled conformation in trajectories of the mutant HRAS. As to the target states, we described three such scenarios as shown in Figure 8. The target state-1 was described, as the open state of Switch I and the closed state of Switch II, whereas the target state-2 was described as the partially open state of Switch I and the open state of Switch II. The target state-3 corresponded to the open state of Switch I and II as shown in Figure 8. We applied the PSP method ***Jalalypour et al. (2020***) on the three scenarios given in Figure 8.

**Figure 8.**
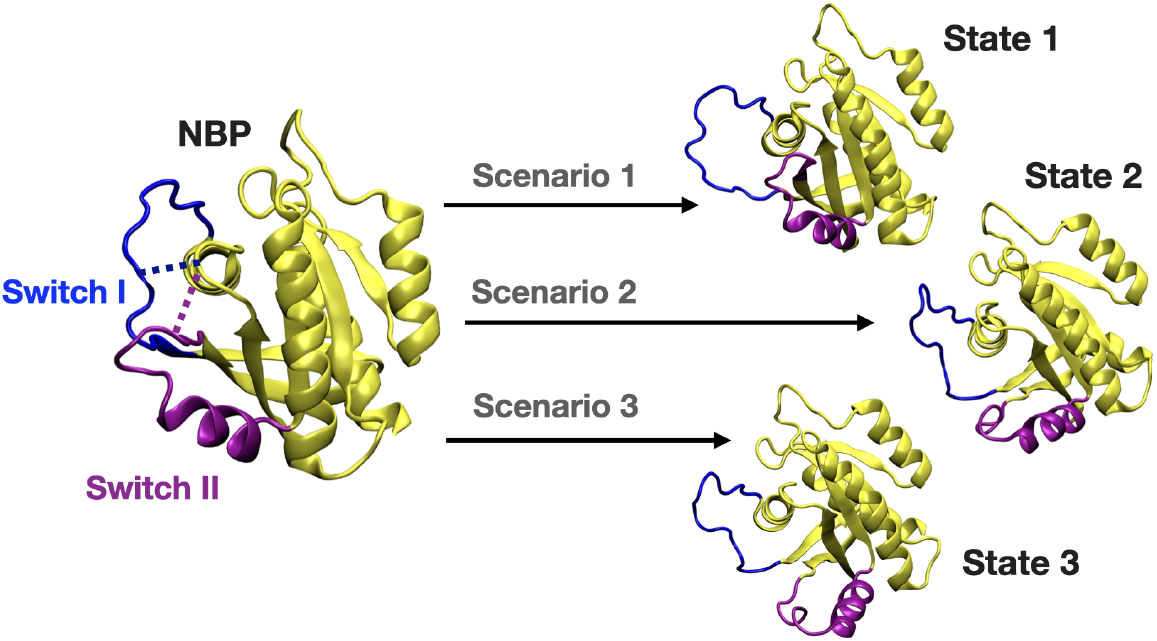
A schematic that illustrates the PRS calculations made for examining the transition between the initial and target states. The initial state represents the conformation of the closed-state of Switch I and II. The target state-1 is described as the open state of Switch I (blue) and close state of Switch II (purple). The target state-2 represents the partially open state of Switch I and open state of Switch II. The target state-3 corresponds to open state of Switch I and II.

The results showed that transition between the initial state and the target state-1 gave the highest overlap compared to other two states of the final state as shown in Table 3. Therefore, this finding shows that Switch I residues mainly contributed to the conformational transition of displacement of Switch I out of the nucleotide-binding pocket in HRAS^G12D^.

**Table 3.** The results of PRS calculations for the transition between initial and target states.

| Ligand | State | D12-P34 (Å) | G60-GTP (Å) | PRS selected residues | PRS overlap(*O*<sup>i</sup>) |
| --- | --- | --- | --- | --- | --- |
| Cerubidine<sup>a</sup> | Initial state<sup>b</sup> | 7.7 (closed) | 7.0 (closed) | - | - |
|  | **Target state-1** | **26.3 (open)** | **11.00 (closed)** | **34, 35, 33, 32, 37, 36** | **0.74-0.70** |
|  | Target state-2 | 13.9 (partially open) | 15.00 (open) | 35, 34, 33, 66, 16, 65 | 0.58-0.50 |
|  | Target state-3 | 19.5 (open) | 16.9 (open) | 34, 66, 35, 64, 16, 33 | 0.59-0.50 |

We further investigated if cerubidine facilitated displacement of Switch I in terms of energetic cost required. To this end, we performed steered molecular dynamics simulations by using ligandfree and cerubidine-bound HRAS^G12D^ systems using the coordinates obtained by PRS method as shown in bold in Table 3. In particular, Y32 and its best direction with overlap values (*O*^*i*^) of 0.72 were fed to SMD simulation. The initial structure was then perturbed by pulling the C_*α*_ atom of Y32 along the best direction towards the target state-1. Each simulation was repeated ten times and the potential of mean force (PMF) was calculated. Results indicated significant energetic difference between PMF profiles pertaining to ligand-free and cerubidine-bound systems (Figure 9). Consequently, this finding showed that cerubidine facilitated opening of the Switch I and exposure of Y32.

**Figure 9.**
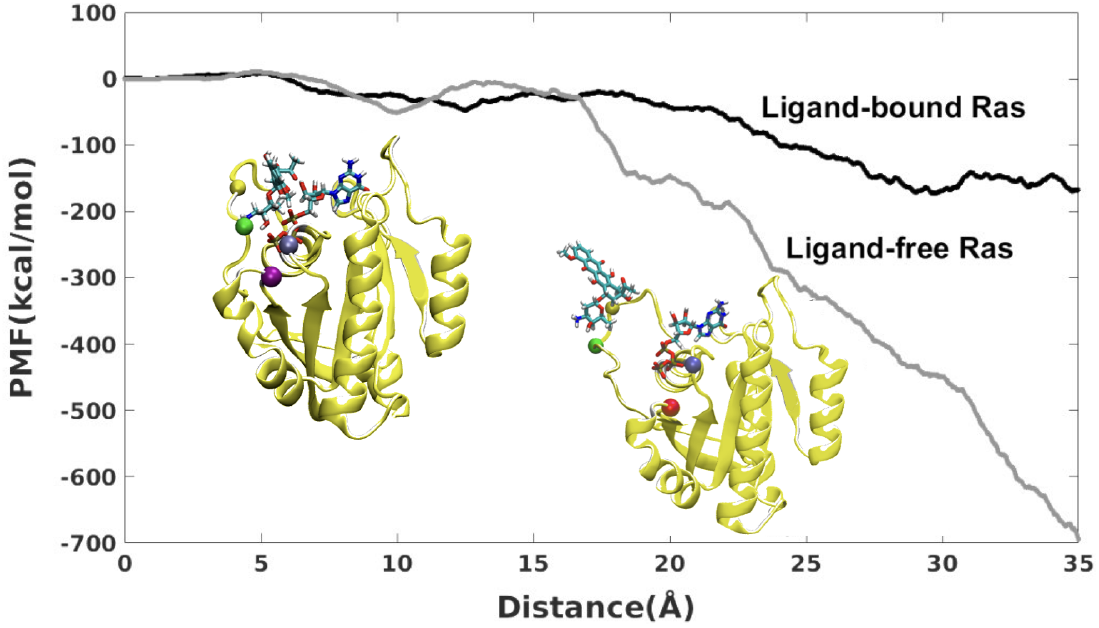
PMF along the PSP predicted coordinate with the highest overlap for the transition scenarios 1 (Switch I opening motion) as a function of distance. PMF is calculated for HRAS^G12D^ system in the presence and absence of cerubidine, and each simulation was repeated ten times. The distance was calculated between the initial and final position of the SMD atom (shown as a yellow bead). The initial and final structures of an SMD simulation were illustrated on the left and right sides of the figure, respectively. Yellow bead: Y32; Green bead: P34; Iceblue bead: G60; GTP and Cerubidine: Licorice representation; Green line: The distance between D12 and P34; Purple line: The distance between G60 and GTP.

### HEK-293T cells were engineered to express HRAS^G12D^ mutant

To investigate G12D specific system properties *in vitro*, we established G12D mutant HRAS expressing cell lines. Human embryonic kidney cells (HEK-293T; CRL-11268, ATCC) is a widely used cell line for gene delivery studies due to their high transfection efficiencies ***Ooi et al. (2016***). Accordingly, as proof of concept, we aimed to introduce G12D mutant HRAS into HEK-293T cells (293T-HRAS^G12D^). We firstly subcloned the HRAS^G12D^ gene region in the commercially available plasmid with a bacterial expression system into the eukaryotic expression plasmid carrying a puroR gene as a selection marker (See Figure 10.A). Next, we showed that the transfection method reaches a high efficiency (90-95%) when a GFP expressing plasmid is introduced into HEK-293T cells (See Figure 10.B).

**Figure 10.**
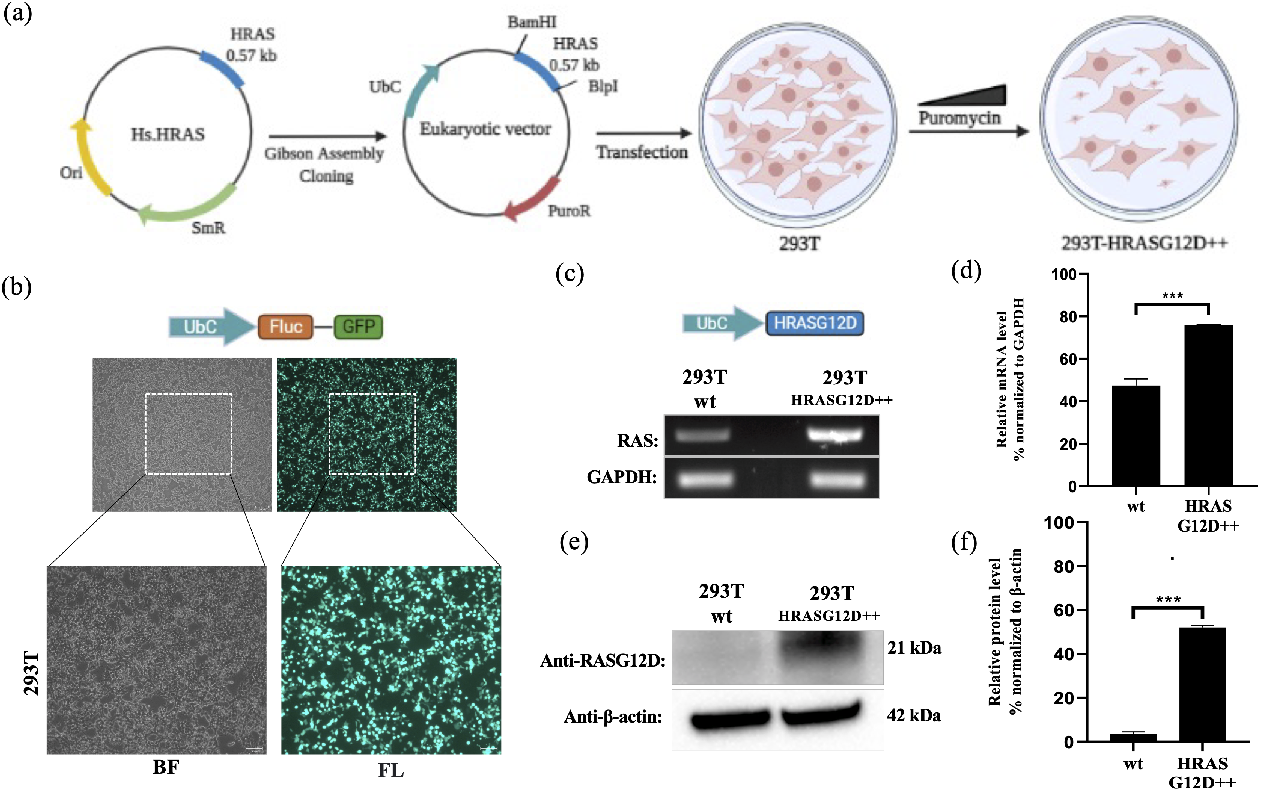
Engineering HEK-293T cells expressing mutant HRAS^G12D^. (a) Schematic representation of the cloning HRAS^G12D^ gene region into the eukaryotic expression plasmid (with PuroR gene to select transgene positive population) using the Gibson Assembly method and engineering HEK-293T cell line to overexpress HRAS^G12D^ protein upon transfection followed by puromycin selection. (b) Fluorescent images 293T cells transfected with GFP-encoding plasmid. (c) RT-PCR analysis showing expression levels of HRAS^G12D^ in 293T cells transfected with HRAS^G12D^ plasmid. (d) ImageJ analysis of band densities from “C”. (e) Western blot analysis showing expression levels of HRAS^G12D^ in 293T-HRAS^G12D^ cells. (f) ImageJ analysis of Western-blot band densities. Data represent the means of three independent assays. Unpaired t-test analysis was used to test the difference between each experimental group and the control group. BF: bright field, FL: Fluorescence, ***: p<0,0001

A cDNA library of the 293T-HRAS^G12D^ cell lysates was obtained for RT-PCR analysis and we detected that the 293T-HRAS^G12D^ cells express increased levels of HRAS transcripts compared to control 293T cells (wild-type; cells with no gene transfer) (See Figure 10.C and D). The primer sets can amplify both wild-type and mutant forms of HRAS since there is only one single base difference and no mutant specificity ***Muñoz-Maldonado et al. (2019***). Therefore, we detected HRAS^G12D^ expression at the protein level using a G12D specific antibody. Our results showed that the transfected cells (293T-HRAS^G12D^) express significantly high levels of HRAS^G12D^ protein compared to wild-type cells (See Figure 10.E and F). Interestingly, we found out that wild-type HEK-293T cells naturally lack G12D mutant protein expressions.

### Cerubidine treatment selectively inhibits the HRAS^G12D^-RAF interaction and blocks activation of HRAS^G12D^

To study the potential HRAS^G12D^-RAF targeting effects of our proposed small molecule cerubidine, firstly, we determined the optimum doses of the molecule in 293T cell lines. The cells treated with a range of compound concentrations (1, 5, 10, 25, 50, 100 *μ*M) showed 80% viability up to 10 *μ*M treatment. Besides that, 25 *μ*M and above cerubidine treatments were cytotoxic to the cells (See Figure 11). We then used active RAS pull-down and detection kit (Thermo) to analyze the interaction of the active RAS protein with RAF protein in the presence of cerubidine. To confirm the proper functioning of the kit, we treated 293T cell lysates with GTP*γ*S and GDP *in-vitro* to activate and inactivate RAS. GTP*γ*S is the non-hydrolyzable or slowly hydrolyzable analog of GTP. RAS is active when interacting with GTP and inactive upon binding of GDP ***Simanshu et al. (2017***).In this context, GTP*γ*S was treated with RAS, which increased the interaction of the RAS protein with RAF by keeping it in its active form (See Figure 12.A, B, and C). Following detection of the RAS-RAF interaction, we treated

**Figure 11.**
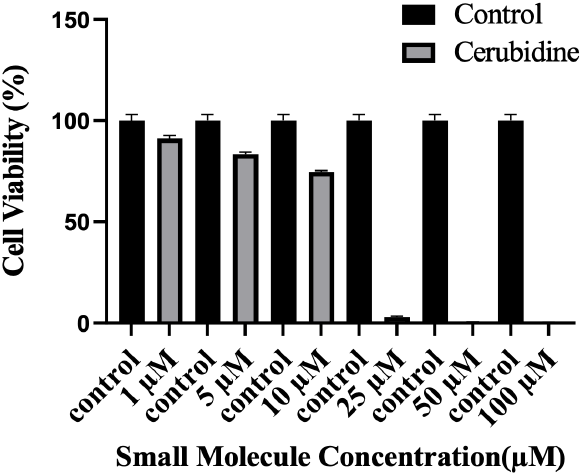
Cell viability assay in control and 24 hr treated 293T cells. The plot indicates the cell viability of 293T cells upon treatment with cerubidine in a dose-dependent manner (1, 5, 10, 25, 50, 100 *μ*M).

**Figure 12.**
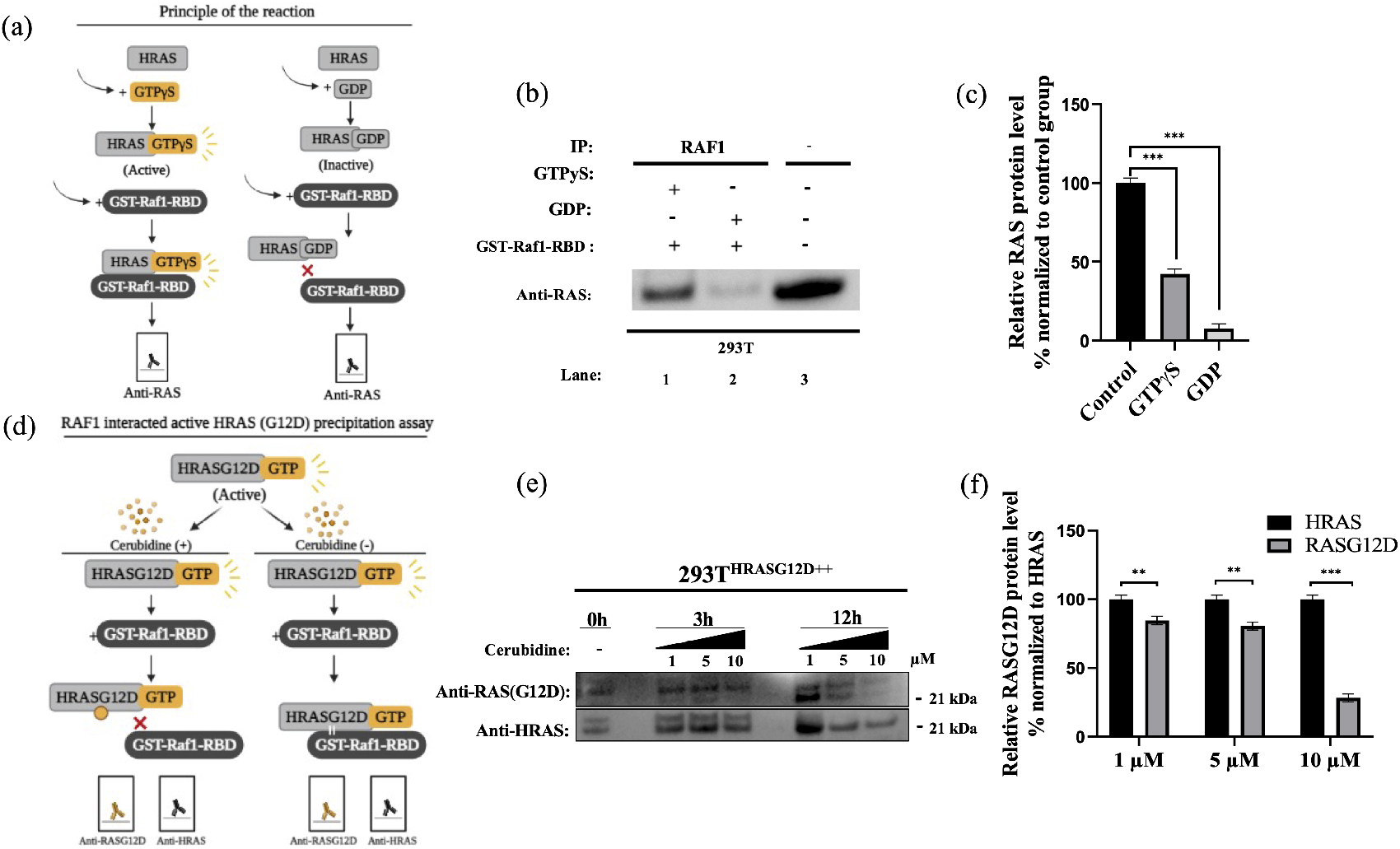
Cell viability assay in control and 24 hr treated 293T cells. (a) Scheme to outline the principle of active Ras pull-down reaction. (b) Immunoprecipitation (IP) assays show interactions of RAS with RAF proteins in the presence (Lane 1-2) and absence (Lane 3) of GTP*γ*S-GDP. Protein extracts were immunoprecipitated with Raf1-RBD probe and resolved by SDS PAGE. Protein-protein interactions were immunodetected using anti-RAS antibodies. (c) ImageJ analysis of Western-blot band densities. Unpaired t-test analysis was used to test the difference between each experimental group and the control group. (d) Scheme outlining the RAF1 interacted active HRAS^G12D^ precipitation assay. (e) Immunoprecipitation (IP) assays showing interactions of RAS with RAF proteins in 293T^HRASG12D++^ cells treated with increasing doses (1,5 and 10 *μ*M) of Cerubudine. Protein extracts obtained at different time points (0h, 3h, and 12h) were immunoprecipitated with the RAF1-RBD probe and resolved by SDS-PAGE. Protein-protein interactions were immunodetected using anti-RAS^G12D^ and anti-HRAS antibodies (f) ImageJ analysis of Western-blot band densities. Unpaired t-test analysis was used to test the difference between each RAS^G12D^ group and the HRAS group. **: p<0.001, ***: p<0.0001).

293T-HRAS^G12D^ cells with optimum doses of cerubidine and collected lysate for protein isolation at different time points. We detected a significant decrease in the active RAS^G12D^, especially at the 12^th^ hr of treatment. Additionally, we analyzed the presence/decrease of active wild-type HRAS in the same line and there was no significant change in active HRAS levels after cerubidine treatment. Overall data showed that the cerubidine treatment blocks HRAS-RAF interaction in a G12D specific manner (See Figure 12.D,E, and F).

## Discussion

Due to involvement in crucial biological processes such as cell growth, proliferation, and differentiation, the RAS protein family has been used as a hot target in drug discovery studies. However, no therapeutic molecule has yet been proven to be used in the clinics due to the absence of deep clefts on the surface of the protein. On the other hand, recently, phosphorylation has been shown to impact the function of the RAS by inhibiting its interaction with effector proteins like RAF, which is involved in the onset of various cancer types. Moreover, examination of the crystal structures pertaining to RAS/RAF complexes showed that Y32 was pointing towards the nucleotide-binding pocket, whereas it was far in the RAF inhibitor-bound RAS protein (PDB ID: 6WGN) ***Zhang et al. (2020***) suggesting that orientational preference of Y32 might control interaction of RAS with RAF (See Figure 13).

**Figure 13.**
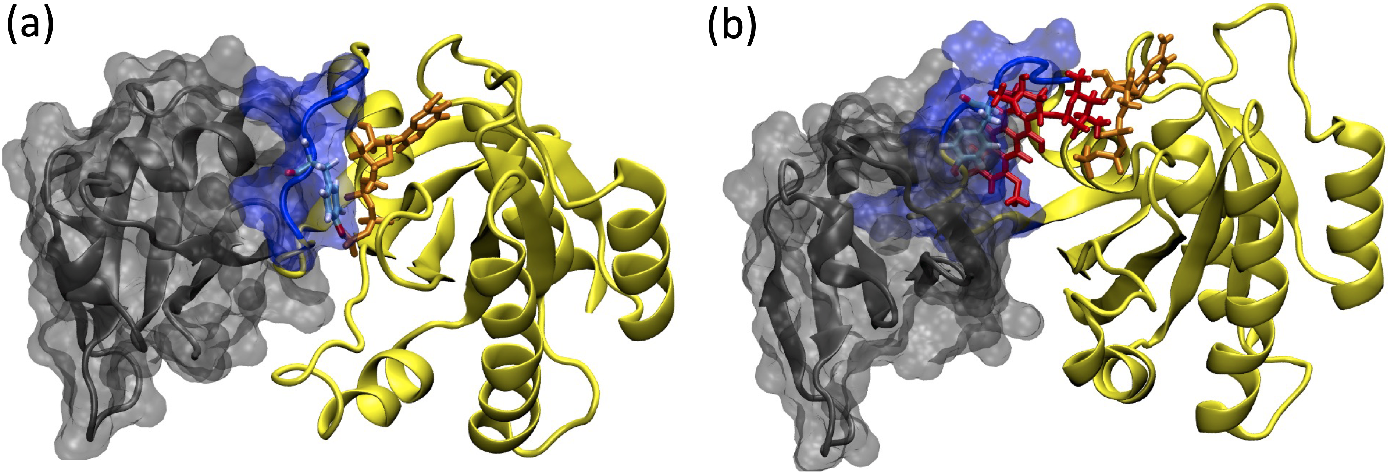
(a)RAF-RBD in complex with HRAS. Y32 and GTP are shown in licorice representation, whereas protein and RAF-RBD interaction interface is shown in New Cartoon, and surface representation, respectively. (b) The displacement of Y32 from the nucleotide-binding pocket by cerubidine, which is colored with red, causes steric clash at the RAS/RAF interface

In this study, motivated by these structural and biochemical data we set out to investigate the impact of phosphorylation on dynamics and structure of HRAS^WT^, and aimed to induce similar modulation in the mutant HRAS by means of small therapeutics to prevent interaction with RAF. To this end, we performed extensive MD simulations on the phosphorylated HRAS and showed that the post-translational modification impacted the dynamics of Switch I and also pushed Y32 out of the nucleotide-binding pocket. Importantly, flexibility of Switch I in the mutant RAS provided a possible binding pocket in the vicinity of the nucleotide-binding site which could be targeted by FDA-approved ligands that modulated dynamics of Y32. Moreover, we also showed that displacement of Switch I and Y32 by the ligand was energetically more favorable than in the absence of the ligand.

Cancer cells show highly mutagenic profile and hard to treat with standard therapies without cancer cell selectivity. Additionaly, in today’s medicine, personalized approaches are ultimately needed considering the individual based differences of the pathology. HRAS mutations are very common in cancer and G12D variant is primarily found in bladder urothelial carcinoma, cutaneous melanoma, infiltrating renal pelvis, ureter urothelial carcinoma, melanoma, and colorectal adenocarcinoma ***Consortium (2017***). Accordingly, in our study, we showed that cancer specific G12D mutant can be targeted by small molecules to interfere with RAF interaction and eventually RAS inactivation. Targeting HRAS-G12D by small molecules can be adopted to further study cell proliferation/death kinetics considering inhibition of RAF/MEK/ERK signalling. Here, we studied HRAS^G12D^; however, high sequence conservation and phosphorylation present among RAS isoforms suggest the potential application of the methodology to other members in the RAS protein family. From that perspective, this study does not only provide mechanistic insight into the impact of phosphorylation but also opens up new avenues for possible use of the post-translational modification in future drug discovery studies. Hereby, we suggest further preclinical examination of our hypothesis for biological mechanisms which might potentiate their clinical uses.

## Methods and Materials

### Molecular dynamics simulations of GTP-bound HRAS^pY32^

#### System setup for molecular dynamics simulations

The crystal structure of phosphoaminophosphonic acid-guanylate ester (GNP)-bound HRAS^WT^ (PDB ID: 5P21) ***Pai et al. (1990***) was retrieved from the Protein Data Bank (https://www.rcsb.org/) ***Berman et al. (2000); Burley et al. (2019***). In order to prepare its GTP-bound state, the N_3_B atom of GNP was substituted with oxygen atom. The crystal waters, which were located within 5 Å of the nucleotide, were kept in simulations. Following, the GTP-bound form of the protein was protonated at pH 7.4 according to the pKa values obtained from the ProPka server ***Søndergaard et al. (2011); Olsson et al. (2011***). The phosphorylation of Y32 residue was made using the TP2 patch provided by CHARMM-GUI server ***Johnson and Lewis (2001***). The protein, GTP and Mg^2+^ ion were parametrized using the CHARMM36 force-field ***Best et al. (2012***) while water molecules were modeled using the TIP3P water model ***Mark and Nilsson (2001***). The thickness of the water layer was set to 15 Å to take periodic boundary conditions into account. Eventually, the solvated system was neutralized with 0.15 M NaCl.

#### Simulation protocol

The MD simulations were employed via Compute Unified Device Architecture version of Nano-Scale Molecular Dynamics ***Vanommeslaeghe et al. (2010); Best et al. (2012); Vanommeslaeghe and MacKerell Jr (2012); Vanommeslaeghe et al. (2012); Yu et al. (2012); Gutiérrez et al. (2016***), in which the graphical processing unit acceleration was enabled. Temperature, pressure, and time step were set to 310 K, 1 atm, 2 femtoseconds, respectively. In order to calculate the long-range electrostatic interactions, the particle mesh Ewald method was used ***Darden et al. (1993); Essmann et al. (1995***). For computation of non-bonded interactions, the cut-off value was adjusted to 12 Å. Moreover, the prepared system was minimized for 2400 time steps. After minimization, the GTP-bound HRAS^pY32^ system was simulated in the NPT ensemble for a total of *ca*. 2.5 *μ*s. Two simulations were performed each of which started with a different velocity distribution. Obtained trajectories were analyzed by combining these two replicates.

### Ensemble-based virtual screening

#### Clustering the trajectory pertaining to HRAS^G12D^, identification of possible binding pockets, and determination of pharmacophore groups

The most probable conformational state of the binding pocket pertaining to HRAS^G12D^ was determined by using following reaction coordinates: distance measured between i)side-chain oxygen of residue T35 and P*γ* atom of GTP, ii) backbone amide of the residue G60 and P*γ* atom of GTP, and iii) side-chain oxygen of the residue Q61 and P*γ* atom of GTP, which were used in our previous study. The frames, which represent different conformational states of the nucleotide-binding pocket with respect to the above-mentioned coordinates, were selected. Subsequently, GTP and Mg^2+^ were removed from the frames and proteins were optimized using the OPLS3e force-field ***Roos et al. (2019***) that is available in the “Protein Preparation” module of the Schrödinger software ***Sastry et al. (2013); Release (2018); Roos et al. (2019***). The optimized structures were provided as inputs to the “SiteMap” module of the Schrödinger ***Halgren (2007***, 2009); ***Release (2018***). Subsequently, possible binding pockets having higher scores were identified and utilized in further steps. Afterwards, pharmacophore groups were built up in accordance with chemical and geometrical properties of the identified binding pockets. To do so, the “Develop Pharmacophore Model” module of Schrödinger was utilized ***Salam et al. (2009); Loving et al. (2009***). Following, candidate molecules, which include at least 3 of the 7 pharmacophore features and have molecular weight less than 550 kDa, were sought in the BindingDB ***Gilson et al. (2016); Liu et al. (2007); Chen et al. (2001b***, 2002, 2001a), DrugCentral ***Ursu et al. (2016***, 2019), NCGC ***Huang et al. (2011***), and DrugBank ***Wishart et al. (2018); Law et al. (2014); Knox et al. (2010); Wishart et al. (2008***, 2006) databases.

### Testing the stability of ligand-HRAS^G12D^ complexes via atomistic simulations

After the selection of candidates based on their GScore values and orientations next to the nucleotidebinding pocket, the stability and the impact of the ligands on the structure and dynamics of HRAS^G12D^ were explored by means of MD simulations. To this end, the topology and parameter files of the candidate molecules were prepared using the “Ligand Reader & Modeler” of CHARMM-GUI ***Jo et al. (2008); Kim et al. (2017***). The systems were simulated using at least two replicates, each of which started with different initial velocity distribution under the same conditions that were used for HRAS^pY32^. Eventually, ligand-protein complexes were simulated for a total of *ca*. 9.4 *μ*s (See Table S2).

### Local and global analysis of the trajectories

The trajectories were visualized with the “Visual Molecular Dynamics” (VMD) and snapshots were rendered using the “Taychon Render” ***Humphrey et al. (1996); Stone (1998***). “Groningen Machine for Chemical Simulations” (GROMACS) package and ProDy library were utilized for the local and global trajectory analysis ***Abraham et al. (2015); Lindahl et al. (2021); Bakan et al. (2011***).

#### Root-mean-square fluctuation

The root-mean-square fluctuation (RMSF) of backbone atoms throughout the obtained trajectories was calculated using the “gmx rmsf” module of GROMACS ***Abraham et al. (2015); Lindahl et al. (2021***) as shown in the below formula;

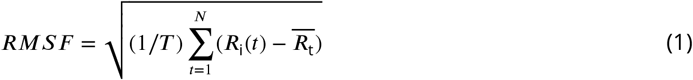

where *T* and *R*_*i*_*(t)* correspond to the duration of simulation and coordinates of backbone atom *R*_*i*_ at time *t*, respectively. By courtesy of this, the flexibility of each residue was computed, and made a holistic comparison with the systems. Particularly, for uncovering the impact of ligands on the backbone RMSF of Y32 and RAF interaction interfaces, the backbone RMSF value of the regions/residues of interest pertaining to ligand-bound HRAS^G12D^ was subtracted from those of HRAS^G12D^.

#### Probability distributions of atom-pair distances

The probability distributions of atom-pair distances were exploited to have a closer look into the impact of the tyrosyl phosphorylation on HRAS^WT^ as well as the impact of candidate molecules on HRAS^G12D^. To this end, the “gmx distance” module of GROMACS was utilized for measuring the distance (i) between the C_*α*_ atoms of G/D12 and P34, and (ii) between the side-chain oxygen atom of Y32 and P*γ* atom of GTP ***Abraham et al. (2015); Lindahl et al. (2021***). The computed raw-data was converted into probability plots by calculating the frequencies of the sampled distances adjusting the sampling range as 2 Å.

### Number of water molecules

The number of water molecules around the GTP was calculated over the course of produced trajectories to reveal the impact of mutation, tyrosyl phosphorylation, and ligands on the exposure of GTP to the possible nucleophilic water attacks via the ProDy library ***Bakan et al. (2011***). To this end, the water molecules within 5 Å of GTP were selected and computed per frame. Thereafter, the mean of the number of water molecules around GTP was taken as well as the standard error of the mean was calculated.

### Principal component analysis

In addition to the local dynamics and structural properties of the phosphorylated system, its overall dynamics were also scrutinized via principal component analysis (PCA). The principal components of HRAS^pY32^ were compared with those of HRAS^WT^, and HRAS^G12D^. By doing so, the collective effect of the tyrosyl phosphorylation was demystified. In this regard, the trajectory of HRAS^pY32^ was aligned with respect to the C_*α*_ atoms of the reference structure, and subsequently, a diagonalized co-variance matrix was generated;

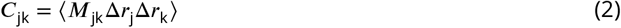

where *M*_*jk*_Δ *r*_*j*_ Δ *r*_*k*_ corresponds to displacement from time-averaged structure for each coordinate of *j* and *k* atoms, whilst co-variance matrix is abbreviated by *C*_*jk*_.

Following the generation of the diagonalized co-variance matrix, eigenvectors (*v*) and eigenvalues (δ^*2*^) were calculated.

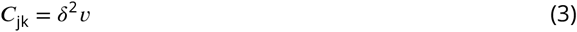

The diagonalized co-variance matrix was generated using the “gmx covar” module of GROMACS ***Abraham et al. (2015); Lindahl et al. (2021***). Thereafter, the “gmx anaeig” module of GROMACS was made use of taking the projection of the trajectory with respect to the eigenvectors of interest, which eventually illuminated the collective spatial organization of the protein as well as the eigen RMSF values of the C_*α*_ atoms ***Abraham et al. (2015); Lindahl et al. (2021***).

### Perturb-scan-pull

PSP consists of three parts, which are PRS, steered molecular dynamics (SMD), and potential of mean force (PMF) calculation ***Jalalypour et al. (2020***). Firstly, the PRS calculation of all the ligand-bound systems were conducted, whilst the SMD and PMF calculation were carried out for the cerubidine-bound HRAS^G12D^ system, whose dynamic and structural properties are similar to those of other studied ligand-bound systems as elucidated by atomistic simulations.

#### Perturbation-response scanning

PRS was performed to achieve the target states by perturbing each residues on the initial state, which, in turn, provided insight into the response of all residues in the HRAS^G12D^. In this way, the residues, which play a pivotal role in the anticipated transitions, were aimed to be identified. To this end, the spatial position of both Switch I and II was clustered to determine initial and target states by measuring the distance between (i)the C_*α*_ atoms of D12 and P34 and (ii) the backbone amide of G60 and P*γ* of GTP over the course of trajectories pertaining to HRAS^G12D^ and ligand-bound HRAS^G12D^. Following, the coarse-grain representation of each state was modeled by selecting the center of mass of the C_*α*_ atom pertaining to each residue as a node. Herein, 1000 random forces (Δ**F**) in distinct directions were sequentially exerted on each node in order to perturb the initial structure ***Atilgan and Atilgan (2009***). In light of the linear response theory, displacement (Δ**R**) as a response to force exerted on the structure was derived from an equilibrated chunk of MD simulations;

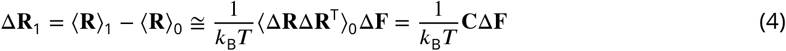

where **R**_**0**_ and **R**_**1**_ correspond to the unperturbed initial state of HRAS^G12D^ and perturbed predicted coordinates, respectively;

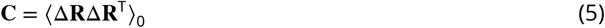

where the cross-correlation of the fluctuations of the nodes in the initial state is denoted by **C**.

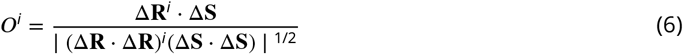

The measured difference between the initial and target structures and the overlap between two nodes are denoted by Δ**S** and *O*^*i*^, respectively.

#### Steered molecular dynamics

Following the PRS calculation, SMD simulations were employed under the same circumstances as the above-mentioned MD simulations pertaining to HRAS^pY32^. The set of external poses were imposed to the C_*α*_ atom of Y32, where the constant velocity and spring constant were adjusted to

0.03 Å ps^-1^ and 90 kcal mol^-1^Å^-2^, respectively. Moreover, the C_*α*_ atoms of L23 and R149 residues were fixed along the pulling direction so as to prevent dislocation and rotation on the structure. The SMD runs were considered completed as long as the secondary structure of the protein was maintained and the final structure resembled the target conformation.

#### Potential of mean force

The energy landscape of the transition in either presence or lack of the drug molecule, namely cerubudine, was elaborated by calculating the PMF along the pulling direction. Considering the well-established procedure ***Jalalypour et al. (2020***), the PMF was computed according to the secondorder cumulant expansion formula via,

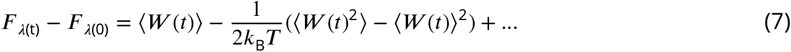

### Preparation of plasmid constructs encoding HRAS^G12D^

The bacterial expression plasmid Hs.HRAS^G12D^ (83183) was obtained from Addgene (U.S). The mutant HRAS^G12D^ was then inserted between the BamHI and BlpI restriction sites under the (UbC promoter) into the lentiviral vectors with the PuroR gene. (The vector was a kind gift from Dr. Shah (Brigham and Woman’s Hospital, Harvard Medical School, Boston, U.S.) and was previously characterized and widely studied ***Stuckey et al. (2015***)).

### Engineering HRAS^G12D^ expressing HEK-293T cells; 293T-HRAS^G12D^

To investigate the *in vitro* outcomes of our *in silico* findings, HEK-293T cells (CRL-11268, ATCC) were engineered to express mutant HRAS^G12D^. HEK-293T cell lines cultured on T75 flask with high glucose DMEM medium which contains 10% fetal bovine serum (FBS), 1% Penicillin Streptomycin at 37 °C in 5% CO_2_ incubator. One day (18 to 24 hr) prior to transfection, cells were seeded at an optimum density that reaches 70-80% confluency the next day, at the time of transfection. Plasmid DNA was transfected into cells using Trans-Hi™ *In Vitro* DNA Transfection Reagent (F90101TH, FormuMax) according to the manufacturer’s recommendations. 12 to 18 hr after transfection, the medium containing the Trans-Hi™/DNA complex was removed and replaced with a fresh whole serum/antibiotic-containing medium.

### HRAS^G12D^ expression analysis

To use 293T-HRAS^G12D^ cells in the following experiments, first of all, we analyzed the overexpression of HRAS transcripts by reverse transcriptase-polymerase chain reaction(RT-PCR). The primer sets can not be specific to the mutant G12D since there is only 1 single base difference in G12D mutant versus wild-type HRAS. For this reason, we further evaluated G12D expression at the protein level via western blot.

#### RT-PCR

To verify increased HRAS transcript levels in HEK-293T cells by RT-PCR, we firstly harvested cells expressing HRAS^WT^ and HRAS^G12D^ to prepare RNA samples. Afterward, RNA was extracted using RNeasy Mini Kit (74104, Qiagen) and the cDNA library was prepared from 1 *μ*g of total RNA, using SuperScript VILO cDNA Synthesis kit (11754050, Invitrogen). HRAS was then amplified by RT-PCR using a standard PCR protocol on a T100 Thermal Cycler (BIO-RAD). Gene expression was normalized to that of a housekeeping gene; GAPDH (glyceraldehyde-3-phosphate dehydrogenase). The primer sets (5’-3’) used for RT-PCR were as follows:

*GAPDH: Fw- GTCAGTGGTGGACCTGACCT; Rv- TGCTGTAGCCAAATTCGTTG* (245bp PCR product) and

*HRAS: Fw- GGATCCATGACGGAATATAAGCTGG; Rv- GCTCAGCTTAGGAGAGCACACACTTGC* (570 bp PCR product)

#### Protein sample preparation

Cells were washed two times with ice-cold phosphate-buffered saline (PBS) prior to 1X lysis buffer (25 mM Tris-HCl,150 mM NaCl, 5 mM MgCl_2_,1%NP-40, and 5% glycerol) involving complete Mini Protease Inhibitor Cocktail tablet (11836153001, Roche). Lysates were spun at 16,000 × g for 15 min and the supernatants were reserved as protein samples. The Pierce BCA Protein Assay Reagent (23227, Thermo Fisher Scientific) was used to quantify the protein concentration of each sample.

#### Western blot (WB) analysis

Cell lysates were resolved by sodium dodecyl sulfate (SDS)-polyacrylamide gel electrophoresis (PAGE) using Bolt™ 4 to 12%, Bis-Tris, 1.0 mm, Mini Protein gel (NW04120BOX, Invitrogen). Proteins were transferred into nitrocellulose membrane by iBlot 2 Dry Blotting System (Invitrogen) at constant current of 1.3 A for 7 min. Membranes were blocked with 5% BSA-ALBUMIN in Trisbuffered saline/0.1% Tween-20 for 1 hr at room temperature and incubated overnight with rabbit anti-RAS^G12D^ (mutant specific) (14429, Cell Signaling Technology) antibody. After primary antibody incubation, membranes were washed with TBST (Tris-buffered saline with 0.05% Tween-20). Secondary antibody (R-05071-500, Advansta), HRP conjugated goat anti-rabbit was diluted to 1:3000 in 5% BSA and incubated for 1 hr at room temperature. Membranes were developed using ECL substrate (1705061, Bio-Rad) and a chemiluminescence signal was detected by Chemidoc (Bio-Rad). Next, *β*-actin levels were determined as loading controls. For this, the membrane was incubated in stripping buffer (0.2 M Glycine, 0.10% Tween-20, pH:2.5) and blocking solution before reprobing with anti-*β*-actin (3700, Cell Signaling Technology).

### Optimization of optimum cerubidine treatment doses to target HRAS^G12D^-RAF interaction

HEK-293T cells were plated at 5000 cells/well into 96 black well plates (3603, Corning) and cultured in DMEM, high glucose (Gibco) containing 10% FBS at 37 °C in 5% CO2. Cells were cultured overnight and the compounds (dissolved in DMSO) were added to the cells at concentrations ranging from 0 to 100 *μ*M. The cells were incubated under standard culture conditions for 24 hr. Cell viability was quantified using the CellTiterGlo Luminescent Cell Viability Assay (Promega) according to the manufacturer’s instructions to measure ATP generated by metabolically active cells. Luminescent signals were measured using the SpectraMAX (Molecular Devices). The luminescence signals obtained from the compound-treated cells were normalized against the signal for DMSO-only treated cells.

### Active RAS pull-down assay

In this experimental setup, we conceptually investigated “G12D versus wild-type” HRAS presence in the active RAS population in the cells treated with cerubidine. RAS activity was determined using Active RAS Pull-Down and Detection Kit (Thermo Fisher Scientific) following the manufacturer’s instructions. Firstly, we tested the assay validity using provided supplements. Lysates were incubated with glutathione S-transferase fusion of the RAS binding domain (RBD) of RAF1 along with glutathione agarose for 1 hr. Agarose beads were collected by centrifugation and washed three times with 1X Wash Buffer (25 mM Tris-HCl,150 mM NaCl, 5 mM MgCl_2_,1%NP-40, and 5%glycerol). Each sample was resuspended and boiled at 100 °C for 5 min. Samples were analyzed by western blotting as previously described. Analysis of RBD pull-down lysates was performed with mouse anti-RAS Antibody (16117, Thermo Fisher Scientific). Secondly, we prepared cell lysates from cerubidine treated 293T-HRAS^G12D^ cells. One day prior to treatment plated a sufficient number of cells so that the cell density reaches the optimal confluency (60-70%) at the time of treatment. Cells were incubated with increased cerubidine concentrations (1, 5, and 10 *μ*M) for 3 hr and 12 hr (0 hr was used as control). After incubation, the active Ras pull-down assay was performed with proteins isolated from the treated and untreated cells (as described above in the protein sample preparation section). Finally, samples were subjected to western blotting as previously described. RBD pull-down lysates were probed with mouse anti-HRAS (sc-29, Santa Cruz Biotechnology), and rabbit anti-RAS^G12D^ Mutant Specific antibodies (14429, Cell Signaling Technology).

## Acknowledgments

MI and OS thank TUBITAK and TUSEB for providing funding in the scope of 2209-A Undergraduate Research Support Program, reference number: 1919B011701434, and Computational Structural Biology Strategic R&D Project Call, reference number: 2019-TA-02-3561, respectively. CA thanks for partial support from TUBITAK, project no. 116F229. MI and OS also acknowledge Istanbul Medipol University for providing access to the High-Performance Computing System so as to run some of the MD simulations. MI, OS, FJ, and CA thank TUBITAK ULAKBIM for giving access to High Performance and Grid Computing Center (TRUBA resources) to complete the rest of the numerical calculations reported in this paper. NK thanks to Dr. Khalid Shah from Brigham and Woman’s Hospital, Harvard Medical School, U.S. for the plasmid backbones.

## References

Abraham MJ, Murtola T, Schulz R, Páll S, Smith JC, Hess B, Lindahl E. GROMACS: High performance molecular simulations through multi-level parallelism from laptops to supercomputers. SoftwareX. 2015; 1:19–25.

Araki M, Shima F, Yoshikawa Y, Muraoka S, Ijiri Y, Nagahara Y, Shirono T, Kataoka T, Tamura A. Solution structure of the state 1 conformer of GTP-bound H-Ras protein and distinct dynamic properties between the state 1 and state 2 conformers. Journal of Biological Chemistry. 2011; 286(45):39644–39653.

Atilgan C, Atilgan AR. Perturbation-response scanning reveals ligand entry-exit mechanisms of ferric binding protein. PLoS Comput Biol. 2009; 5(10):e1000544.

Bakan A, Meireles LM, Bahar I. ProDy: protein dynamics inferred from theory and experiments. Bioinformatics. 2011; 27(11):1575–1577.

Barbacid M. Ras genes. Annual review of biochemistry. 1987; 56(1):779–827.

Berman HM, Westbrook J, Feng Z, Gilliland G, Bhat TN, Weissig H, Shindyalov IN, Bourne PE. The protein data bank. Nucleic acids research. 2000; 28(1):235–242.

Best RB, Zhu X, Shim J, Lopes PE, Mittal J, Feig M, MacKerell Jr AD. Optimization of the additive CHARMM allatom protein force field targeting improved sampling of the backbone Ф, *ψ* and side-chain *χ*1 and *χ*2 dihedral angles. Journal of chemical theory and computation. 2012; 8(9):3257–3273.

Bourne HR, Sanders DA, McCormick F. The GTPase superfamily: conserved structure and molecular mechanism. Nature. 1991; 349(6305):117–127.

Buhrman G, Holzapfel G, Fetics S, Mattos C. Allosteric modulation of Ras positions Q61 for a direct role in catalysis. Proceedings of the National Academy of Sciences. 2010; 107(11):4931–4936.

Bunda S, Heir P, Srikumar T, Cook JD, Burrell K, Kano Y, Lee JE, Zadeh G, Raught B, Ohh M. Src promotes GTPase activity of Ras via tyrosine 32 phosphorylation. Proceedings of the National Academy of Sciences. 2014; 111(36):E3785–E3794.

Burley SK, Berman HM, Bhikadiya C, Bi C, Chen L, Di Costanzo L, Christie C, Dalenberg K, Duarte JM, Dutta S, et al. RCSB Protein Data Bank: biological macromolecular structures enabling research and education in fundamental biology, biomedicine, biotechnology and energy. Nucleic acids research. 2019; 47(D1):D464–D474.

Canon J, Rex K, Saiki AY, Mohr C, Cooke K, Bagal D, Gaida K, Holt T, Knutson CG, Koppada N, et al. The clinical KRAS (G12C) inhibitor AMG 510 drives anti-tumour immunity. Nature. 2019; 575(7781):217–223.

Chen X, Gilson M, Mk B. A Web-Accessible Molecular Recognition Database [Internet]. Comb Chem High Throughput Screen. 2001; p. 719–25.

Chen X, Lin Y, Gilson MK. The binding database: overview and user’s guide. Biopolymers: Original Research on Biomolecules. 2001; 61(2):127–141.

Chen X, Lin Y, Liu M, Gilson MK. The Binding Database: data management and interface design. Bioinformatics. 2002; 18(1):130–139.

Cherfils J, Zeghouf M. Regulation of small gtpases by gefs, gaps, and gdis. Physiological reviews. 2013; 93(1):269–309.

Consortium APG. AACR Project GENIE: powering precision medicine through an international consortium. Cancer discovery. 2017; 7(8):818–831.

Cox AD, Fesik SW, Kimmelman AC, Luo J, Der CJ. Drugging the undruggable RAS: mission possible? Nature reviews Drug discovery. 2014; 13(11):828–851.

Darden T, York D, Pedersen L. Particle mesh Ewald: An N log (N) method for Ewald sums in large systems. The Journal of chemical physics. 1993; 98(12):10089–10092.

De Luca A, Maiello MR, D’Alessio A, Pergameno M, Normanno N. The RAS/RAF/MEK/ERK and the PI3K/AKT signalling pathways: role in cancer pathogenesis and implications for therapeutic approaches. Expert opinion on therapeutic targets. 2012; 16(sup2):S17–S27.

Downward J. The ras superfamily of small GTP-binding proteins. Trends in biochemical sciences. 1990; 15(12):469–472.

Drugan JK, Khosravi-Far R, White MA, Der CJ, Sung YJ, Hwang YW, Campbell SL. Ras Interaction with Two Distinct Binding Domains in Raf-1 5 Be Required for Ras Transformation *. Journal of Biological Chemistry. 1996; 271(1):233–237.

Duffy MJ, Crown J. Drugging “undruggable” genes for cancer treatment: Are we making progress? International Journal of Cancer. 2021; 148(1):8–17.

Eser S, Schnieke A, Schneider G, Saur D. Oncogenic KRAS signalling in pancreatic cancer. British journal of cancer. 2014; 111(5):817–822.

Essmann U, Perera L, Berkowitz ML, Darden T, Lee H, Pedersen LG. A smooth particle mesh Ewald method. The Journal of chemical physics. 1995; 103(19):8577–8593.

Ferro E, Trabalzini L. RalGDS family members couple Ras to Ral signalling and that’s not all. Cellular signalling. 2010; 22(12):1804–1810.

Fetics SK, Guterres H, Kearney BM, Buhrman G, Ma B, Nussinov R, Mattos C. Allosteric effects of the oncogenic RasQ61L mutant on Raf-RBD. Structure. 2015; 23(3):505–516.

Geyer M, Wittinghofer A. GEFs, GAPs, GDIs and effectors: taking a closer (3D) look at the regulation of Rasrelated GTP-binding proteins. Current opinion in structural biology. 1997; 7(6):786–792.

Gilson MK, Liu T, Baitaluk M, Nicola G, Hwang L, Chong J. BindingDB in 2015: a public database for medicinal chemistry, computational chemistry and systems pharmacology. Nucleic acids research. 2016; 44(D1):D1045–D1053.

Grand RJ, Owen D. The biochemistry of ras p21. Biochemical Journal. 1991; 279(3):609–631.

Gutiérrez IS, Lin FY, Vanommeslaeghe K, Lemkul JA, Armacost KA, Brooks III CL, MacKerell Jr AD. Parametrization of halogen bonds in the CHARMM general force field: Improved treatment of ligand–protein interactions. Bioorganic & medicinal chemistry. 2016; 24(20):4812–4825.

Gysin S, Salt M, Young A, McCormick F. Therapeutic strategies for targeting ras proteins. Genes & cancer. 2011; 2(3):359–372.

Hah JH, Zhao M, Pickering CR, Frederick MJ, Andrews GA, Jasser SA, Fooshee DR, Milas ZL, Galer C, Sano D, et al. HRAS mutations and resistance to the epidermal growth factor receptor tyrosine kinase inhibitor erlotinib in head and neck squamous cell carcinoma cells. Head & neck. 2014; 36(11):1547–1554.

Halgren TA. Identifying and characterizing binding sites and assessing druggability. Journal of chemical information and modeling. 2009; 49(2):377–389.

Halgren T. New method for fast and accurate binding-site identification and analysis. Chemical biology & drug design. 2007; 69(2):146–148.

Holderfield M, Deuker MM, McCormick F, McMahon M. Targeting RAF kinases for cancer therapy: BRAF-mutated melanoma and beyond. Nature Reviews Cancer. 2014; 14(7):455–467.

Huang L, Hofer F, Martin GS, Kim SH. Structural basis for the interaction of Ras with RaIGDS. Nature structural biology. 1998; 5(6):422–426.

Huang R, Southall N, Wang Y, Yasgar A, Shinn P, Jadhav A, Nguyen DT, Austin CP. The NCGC pharmaceutical collection: a comprehensive resource of clinically approved drugs enabling repurposing and chemical genomics. Science translational medicine. 2011; 3(80):80ps16–80ps16.

Humphrey W, Dalke A, Schulten K, et al. VMD: visual molecular dynamics. Journal of molecular graphics. 1996; 14(1):33–38.

Ilter M, Sensoy O. Catalytically Competent Non-transforming H-RAS G12P Mutant Provides Insight into Molecular Switch Function and GAP-independent GTPase Activity of RAS. Scientific reports. 2019; 9(1):1–10.

Jalalypour F, Sensoy O, Atilgan C. Perturb–Scan–Pull: A Novel Method Facilitating Conformational Transitions in Proteins. Journal of Chemical Theory and Computation. 2020; 16(6):3825–3841.

Jo S, Kim T, Iyer VG, Im W. CHARMM-GUI: a web-based graphical user interface for CHARMM. Journal of computational chemistry. 2008; 29(11):1859–1865.

Johnson CW, Mattos C. The allosteric switch and conformational states in Ras GTPase affected by small molecules. The Enzymes. 2013; 33:41–67.

Johnson CW, Reid D, Parker JA, Salter S, Knihtila R, Kuzmic P, Mattos C. The small GTPases K-Ras, N-Ras, and H-Ras have distinct biochemical properties determined by allosteric effects. Journal of Biological Chemistry. 2017; 292(31):12981–12993.

Johnson LN, Lewis RJ. Structural basis for control by phosphorylation. Chemical reviews. 2001; 101(8):2209–2242.

Kano Y, Gebregiworgis T, Marshall CB, Radulovich N, Poon BP, St-Germain J, Cook JD, Valencia-Sama I, Grant BM, Herrera SG, et al. Tyrosyl phosphorylation of KRAS stalls GTPase cycle via alteration of switch I and II conformation. Nature communications. 2019; 10(1):1–14.

Khan AQ, Kuttikrishnan S, Siveen KS, Prabhu KS, Shanmugakonar M, Al-Naemi HA, Haris M, Dermime S, Uddin S. RAS-mediated oncogenic signaling pathways in human malignancies. In: Seminars in Cancer Biology, vol. 54 Elsevier; 2019. p. 1–13.

Khan I, MarElia-Bennet C, Lefler J, Zuberi M, Denbaum E, Koide A, Connor DM, Broome AM, Pécot T, Timmers C, et al. Targeting the KRAS *α*4-*α*5 allosteric interface inhibits pancreatic cancer tumorigenesis. Small GTPases. 2021; p. 1–14.

Khan I, Rhett JM, O’Bryan JP. Therapeutic targeting of RAS: new hope for drugging the “undruggable”. Biochimica et Biophysica Acta (BBA)-Molecular Cell Research. 2020; 1867(2):118570.

Kim S, Lee J, Jo S, Brooks III CL, Lee HS, Im W. CHARMM-GUI ligand reader and modeler for CHARMM force field generation of small molecules. Journal of Computational Chemistry. 2017; 38(21):1879–1886.

Knight T, Irving JAE. Ras/Raf/MEK/ERK pathway activation in childhood acute lymphoblastic leukemia and its therapeutic targeting. Frontiers in oncology. 2014; 4:160.

Knox C, Law V, Jewison T, Liu P, Ly S, Frolkis A, Pon A, Banco K, Mak C, Neveu V, et al. DrugBank 3.0: a comprehensive resource for ‘omics’ research on drugs. Nucleic acids research. 2010; 39(suppl_1):D1035–D1041.

Krens LL, Baas JM, Gelderblom H, Guchelaar HJ. Therapeutic modulation of k-ras signaling in colorectal cancer. Drug discovery today. 2010; 15(13-14):502–516.

Lamontanara AJ, Georgeon S, Tria G, Svergun DI, Hantschel O. The SH2 domain of Abl kinases regulates kinase autophosphorylation by controlling activation loop accessibility. Nature communications. 2014; 5(1):1–11.

Law V, Knox C, Djoumbou Y, Jewison T, Guo AC, Liu Y, Maciejewski A, Arndt D, Wilson M, Neveu V, et al. DrugBank 4.0: shedding new light on drug metabolism. Nucleic acids research. 2014; 42(D1):D1091–D1097.

Ledford H. Cancer: the Ras renaissance. Nature News. 2015; 520(7547):278.

Lindahl, Abraham, Hess, van der Spoel, GROMACS 2021 Manual. Zenodo; 2021.

Liu T, Lin Y, Wen X, Jorissen RN, Gilson MK. BindingDB: a web-accessible database of experimentally determined protein–ligand binding affinities. Nucleic acids research. 2007; 35(suppl_1):D198–D201.

Loving K, Salam NK, Sherman W. Energetic analysis of fragment docking and application to structure-based pharmacophore hypothesis generation. Journal of computer-aided molecular design. 2009; 23(8):541–554.

Lowy DR, Zhang K, Declue JE, Willumsen BM. Regulation of p21ras activity. Trends in Genetics. 1991; 7(11-12):346–351.

Lu S, Jang H, Muratcioglu S, Gursoy A, Keskin O, Nussinov R, Zhang J. Ras conformational ensembles, allostery, and signaling. Chemical reviews. 2016; 116(11):6607–6665.

Lu S, Jang H, Zhang J, Nussinov R. Inhibitors of Ras–SOS interactions. ChemMedChem. 2016; 11(8):814–821.

Malumbres M, Barbacid M. RAS oncogenes: the first 30 years. Nature Reviews Cancer. 2003; 3(6):459–465.

Mark P, Nilsson L. Structure and dynamics of the TIP3P, SPC, and SPC/E water models at 298 K. The Journal of Physical Chemistry A. 2001; 105(43):9954–9960.

McCarthy MJ, Pagba CV, Prakash P, Naji AK, van der Hoeven D, Liang H, Gupta AK, Zhou Y, Cho KJ, Hancock JF, et al. Discovery of high-affinity noncovalent allosteric KRAS inhibitors that disrupt effector binding. ACS omega. 2019; 4(2):2921–2930.

McCormick F. KRAS as a therapeutic target. Clinical Cancer Research. 2015; 21(8):1797–1801.

McCormick F. The potential of targeting Ras proteins in lung cancer. Expert opinion on therapeutic targets. 2015; 19(4):451–454.

Milroy LG, Ottmann C. The renaissance of Ras. ACS chemical biology. 2014; 9(11):2447–2458.

Muñoz-Maldonado C, Zimmer Y, Medová M. A comparative analysis of individual RAS mutations in cancer biology. Frontiers in oncology. 2019; 9:1088.

Myers MB, Banda M, McKim KL, Wang Y, Powell MJ, Parsons BL. Breast cancer heterogeneity examined by high-sensitivity quantification of PIK3CA, KRAS, HRAS, and BRAF mutations in normal breast and ductal carcinomas. Neoplasia. 2016; 18(4):253–263.

Olsson MH, Søndergaard CR, Rostkowski M, Jensen JH. PROPKA3: consistent treatment of internal and surface residues in empirical p K a predictions. Journal of chemical theory and computation. 2011; 7(2):525–537.

Ooi A, Wong A, Esau L, Lemtiri-Chlieh F, Gehring C. A guide to transient expression of membrane proteins in HEK-293 cells for functional characterization. Frontiers in physiology. 2016; 7:300.

Ostrem JM, Peters U, Sos ML, Wells JA, Shokat KM. K-Ras (G12C) inhibitors allosterically control GTP affinity and effector interactions. Nature. 2013; 503(7477):548–551.

O’Bryan JP. Pharmacological targeting of RAS: recent success with direct inhibitors. Pharmacological research. 2019; 139:503–511.

Pai EF, Krengel U, Petsko GA, Goody RS, Kabsch W, Wittinghofer A. Refined crystal structure of the triphosphate conformation of H-ras p21 at 1.35 A resolution: implications for the mechanism of GTP hydrolysis. The EMBO journal. 1990; 9(8):2351–2359.

Prior IA, Hood FE, Hartley JL. The frequency of Ras mutations in cancer. Cancer research. 2020; 80(14):2969–2974.

Prior IA, Lewis PD, Mattos C. A comprehensive survey of Ras mutations in cancer. Cancer research. 2012; 72(10):2457–2467.

Release S. 4: Glide. Schrödinger, LLC, New York, NY. 2018;.

Roos K, Wu C, Damm W, Reboul M, Stevenson JM, Lu C, Dahlgren MK, Mondal S, Chen W, Wang L, et al. OPLS3e: Extending force field coverage for drug-like small molecules. Journal of chemical theory and computation. 2019; 15(3):1863–1874.

Ruess DA, Heynen GJ, Ciecielski KJ, Ai J, Berninger A, Kabacaoglu D, Görgülü K, Dantes Z, Wörmann SM, Diakopoulos KN, et al. Mutant KRAS-driven cancers depend on PTPN11/SHP2 phosphatase. Nature medicine. 2018; 24(7):954–960.

Salam NK, Nuti R, Sherman W. Novel method for generating structure-based pharmacophores using energetic analysis. Journal of chemical information and modeling. 2009; 49(10):2356–2368.

Sastry GM, Adzhigirey M, Day T, Annabhimoju R, Sherman W. Protein and ligand preparation: parameters, protocols, and influence on virtual screening enrichments. Journal of computer-aided molecular design. 2013; 27(3):221–234.

Shima F, Ijiri Y, Muraoka S, Liao J, Ye M, Araki M, Matsumoto K, Yamamoto N, Sugimoto T, Yoshikawa Y, et al. Structural basis for conformational dynamics of GTP-bound Ras protein. Journal of Biological Chemistry. 2010; 285(29):22696–22705.

Simanshu DK, Nissley DV, McCormick F. RAS proteins and their regulators in human disease. Cell. 2017; 170(1):17–33.

Søndergaard CR, Olsson MH, Rostkowski M, Jensen JH. Improved treatment of ligands and coupling effects in empirical calculation and rationalization of p K a values. Journal of chemical theory and computation. 2011; 7(7):2284–2295.

Stephen AG, Esposito D, Bagni RK, McCormick F. Dragging ras back in the ring. Cancer cell. 2014; 25(3):272–281.

Stone J, An Efficient Library for Parallel Ray Tracing and Animation; 1998.

Stuckey DW, Hingtgen SD, Karakas N, Rich BE, Shah K. Engineering toxin-resistant therapeutic stem cells to treat brain tumors. Stem cells. 2015; 33(2):589–600.

Takai Y, Sasaki T, Matozaki T. Small GTP-binding proteins. Physiological reviews. 2001; 81(1):153–208.

Travers T, López CA, Van QN, Neale C, Tonelli M, Stephen AG, Gnanakaran S. Molecular recognition of RAS/RAF complex at the membrane: Role of RAF cysteine-rich domain. Scientific reports. 2018; 8(1):1–15.

Ursu O, Holmes J, Bologa CG, Yang JJ, Mathias SL, Stathias V, Nguyen DT, Schürer S, Oprea T. DrugCentral 2018: an update. Nucleic acids research. 2019; 47(D1):D963–D970.

Ursu O, Holmes J, Knockel J, Bologa CG, Yang JJ, Mathias SL, Nelson SJ, Oprea TI. DrugCentral: online drug compendium. Nucleic acids research. 2016; p. gkw993.

Vanommeslaeghe K, Hatcher E, Acharya C, Kundu S, Zhong S, Shim J, Darian E, Guvench O, Lopes P, Vorobyov I, et al. CHARMM general force field: A force field for drug-like molecules compatible with the CHARMM all-atom additive biological force fields. Journal of computational chemistry. 2010; 31(4):671–690.

Vanommeslaeghe K, MacKerell Jr AD. Automation of the CHARMM General Force Field (CGenFF) I: bond perception and atom typing. Journal of chemical information and modeling. 2012; 52(12):3144–3154.

Vanommeslaeghe K, Raman EP, MacKerell Jr AD. Automation of the CHARMM General Force Field (CGenFF) II: assignment of bonded parameters and partial atomic charges. Journal of chemical information and modeling. 2012; 52(12):3155–3168.

Vetter IR, Wittinghofer A. The guanine nucleotide-binding switch in three dimensions. Science. 2001; 294(5545):1299–1304.

Vigil D, Cherfils J, Rossman KL, Der CJ. Ras superfamily GEFs and GAPs: validated and tractable targets for cancer therapy? Nature Reviews Cancer. 2010; 10(12):842–857.

Wang Y, Ji D, Lei C, Chen Y, Qiu Y, Li X, Li M, Ni D, Pu J, Zhang J, et al. Mechanistic insights into the effect of phosphorylation on Ras conformational dynamics and its interactions with cell signaling proteins. Computational and Structural Biotechnology Journal. 2021;.

Wishart DS, Feunang YD, Guo AC, Lo EJ, Marcu A, Grant JR, Sajed T, Johnson D, Li C, Sayeeda Z, et al. DrugBank 5.0: a major update to the DrugBank database for 2018. Nucleic acids research. 2018; 46(D1):D1074–D1082.

Wishart DS, Knox C, Guo AC, Cheng D, Shrivastava S, Tzur D, Gautam B, Hassanali M. DrugBank: a knowledgebase for drugs, drug actions and drug targets. Nucleic acids research. 2008; 36(suppl_1):D901–D906.

Wishart DS, Knox C, Guo AC, Shrivastava S, Hassanali M, Stothard P, Chang Z, Woolsey J. DrugBank: a comprehensive resource for in silico drug discovery and exploration. Nucleic acids research. 2006; 34(suppl_1):D668–D672.

Wittinghofer A, Pal EF. The structure of Ras protein: a model for a universal molecular switch. Trends in biochemical sciences. 1991; 16:382–387.

Wittinghofer A, Scheffzek K, Ahmadian MR. The interaction of Ras with GTPase-activating proteins. FEBS letters. 1997; 410(1):63–67.

Wittinghofer A, Vetter IR. Structure-function relationships of the G domain, a canonical switch motif. Annual review of biochemistry. 2011; 80:943–971.

Young A, Lou D, McCormick F. Oncogenic and wild-type Ras play divergent roles in the regulation of mitogen-activated protein kinase signaling. Cancer discovery. 2013; 3(1):112–123.

Yu W, He X, Vanommeslaeghe K, MacKerell Jr AD. Extension of the CHARMM general force field to sulfonylcontaining compounds and its utility in biomolecular simulations. Journal of computational chemistry. 2012; 33(31):2451–2468.

Zhang Z, Gao R, Hu Q, Peacock H, Peacock DM, Dai S, Shokat KM, Suga H. GTP-state-selective cyclic peptide ligands of K-Ras (G12D) block its interaction with Raf. ACS central science. 2020; 6(10):1753–1761.

