## Supplementary material for "Inhibition of mutant RAS-RAF interaction by mimicking structural and dynamic properties of phosphorylated RAS": SI

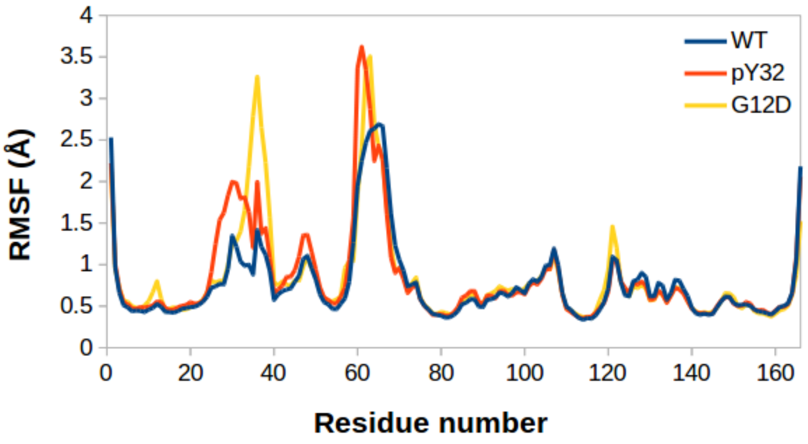


**Figure S1.** The backbone RMSF of H-RAS$^{WT}$, H-RAS$^{pY32}$, and H-RAS$^{G12D}$ was calculated throughout the corresponding trajectories.

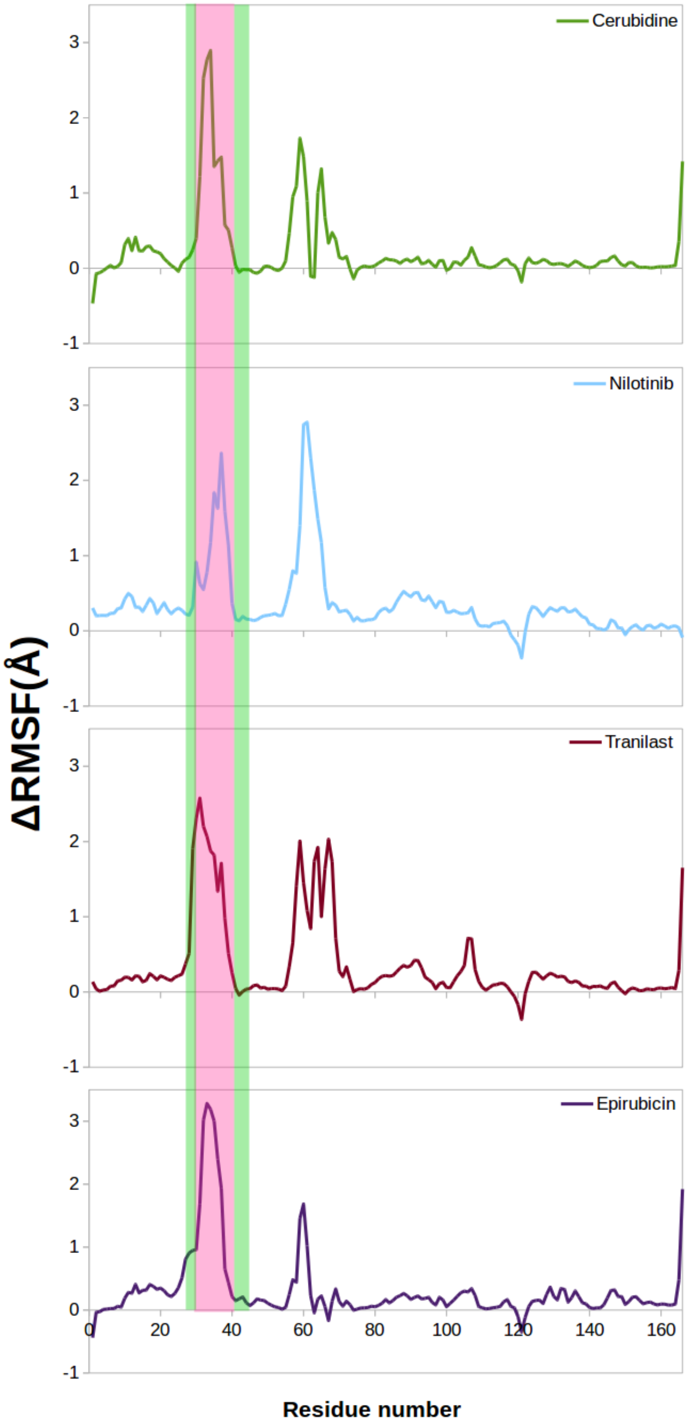


**Figure S2.** The change in the backbone RMSF of H-RAS$^{G12D}$ when the mutant system was treated with cerubidine, nilotinib, tranilast, and epirubicin. The RAF-RBD and -CRD interfaces are shown within the pink and green rectangles.

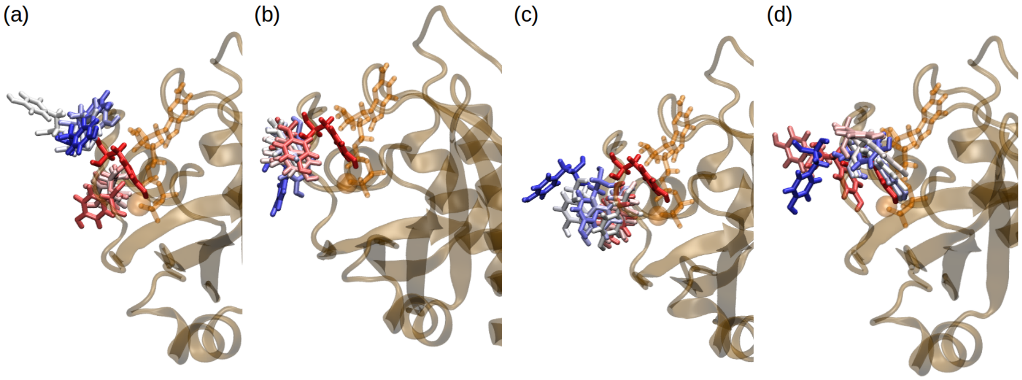


**Figure S3.** The orientational dynamics of Tyr32 over the course of MD trajectories pertaining to the cerubidine-, nilotinib-, tranilast-, and epirubicin-bound H-RAS$^{G12D}$.

**Table S1.** The average number of water molecules within 5 Å of GTP were calculated over the course of the ligand-bound H-RAS$^{G12D}$ systems.

| Ligand-bound H-RAS$^{G12D}$ | $\mu_{water}$ |
|---|---|
| Cerubidine | 118.6±0.1 |
| Tranilast | 110.7 ±0.1 |
| Nilotinib | 86.5 ±0.2 |
| Epirubicin | 104.9 ±0.2 |

**Table S2.** Total simulation time performed for ligand- H-RAS$^{G12D}$ complexes and changes in the backbone RMSF profiles of cerubidine-, tranilast-, nilotinib-, and epirubicin-bound H-RAS$^{G12D}$ systems with respect to those of H-RAS$^{G12D}$.

| Ligand | Duration (ns) | ΔRMSF(Y32) (Å) | $\mu_{\Delta RMSF(RAF-RBD)}$ (Å) | $\mu_{\Delta RMSF(RAF-CRD)}$ (Å) |
|---|---|---|---|---|
| Cerubidine | 1776 | 2.5 | 1.4 | 0.1 |
| Tranilast | 3078 | 2.2 | 1.6 | 0.5 |
| Nilotinib | 1776 | 0.6 | 1.2 | 0.2 |
| Epirubicin | 2736 | 3.0 | 1.9 | 0.5 |

**Table S3.** The results of PRS calculations for the transition between initial and target states.

| Ligand | State | D12-P34 (Å) | G60-GTP (Å) | PRS selected residues | PRS overlap($O^i$) |
|---|---|---|---|---|---|
| Nilotinib | Target state-1 | 22.4 (open) | 5.1 (closed) | 34, 35, 33, 37, 32, 36 | 0.70-0.62 |
| | Target state-2 | 15.3 (partially open) | 12.7 (open) | 61, 62, 63, 23, 22, 65 | 0.57-0.51 |
| | Target state-3 | 22.6 (open) | 19.8 (open) | 34, 35, 33, 37, 36, 66 | 0.66-0.59 |
| Tranilast | Target state-1 | 27.0 (open) | 9.1 (closed) | 32, 36, 33, 37, 34, 35 | 0.74-0.67 |
| | Target state-2 | 14.8 (partially opened) | 12.3 (open) | 22, 18, 23, 104, 87, 6 | 0.55-0.54 |
| | Target state-3 | 20.1 (open) | 14.8 (open) | 34, 33, 35, 32, 37 | 0.54-0.50 |
| Epirubicin | Target state-1 | 29.0 (open) | 7.7 (closed) | 32, 33, 34, 35, 37, 36 | 0.61-0.51 |
| | Target state-2 | 13.3 (partially opened) | 17.8 (open) | 63,62 | 0.51-0.50 |
| | Target state-3 | 22.0 (open) | 15.2 (open) | 34, 35, 33, 37, 36, 66 | 0.61-0.59 |
